# Supercoiling-mediated feedback rapidly couples and tunes transcription

**DOI:** 10.1101/2022.04.20.488937

**Authors:** Christopher P. Johnstone, Kate E. Galloway

## Abstract

Transcription induces a wave of DNA supercoiling, altering the binding affinity of RNA polymerases and reshaping the biochemical landscape of gene regulation. As supercoiling rapidly diffuses, transcription dynamically reshapes the regulation of proximal genes, forming a complex feedback loop. The resulting intergene coupling may provide a mechanism to control transcriptional variance in engineered gene networks and explain the behavior of co-localized native circuits. However, a theoretical framework is needed for integrating both biophysical and biochemical transcriptional regulation to investigate the role of supercoiling-mediated feedback within multi-gene systems. Here, we model transcriptional regulation under the influence of supercoiling-mediated polymerase dynamics, allowing us to identify patterns of expression that result from physical intergene coupling and explore integration of this biophysical model with a set of canonical biochemical gene regulatory systems. We find that gene syntax—the relative ordering and orientation of genes—defines the expression profiles, variance, burst dynamics, and intergene correlation of two-gene systems. By applying our model to both a synthetic toggle switch and the endogenous zebrafish segmentation network, we find that supercoiling can enhance or weaken conventional biochemical regulatory strategies such as mRNA- and protein-mediated feedback loops. Together, our results suggest that supercoiling couples behavior between neighboring genes, representing a novel regulatory mechanism. Integrating biophysical regulation into the analysis and design of gene regulation provides a framework for enhanced understanding of native networks and engineering of synthetic gene circuits.

## 1 Introduction

Cells coordinate complex behaviors through precise spatiotemporal control of gene expression. To rapidly advance gene and cell-based therapies, synthetic biology aims to harness the power of native biology by constructing synthetic gene regulatory networks capable of dynamically prescribing cellular processes, states, and identities [1–4]. Synthetic networks process diverse inputs into complex logical and temporal responses [5–7]. From oscillators to pulse generators, synthetic circuits can precisely coordinate dynamic patterns of gene expression across populations of cells to control cell fate [8–15]. However, rational *de novo* design of synthetic circuits remains challenging. Despite extensive biomolecular modeling, integration of single genetic elements into systems often leads to emergent behaviors, requiring iterative design-build-test cycles to achieve the desired performance [16–18]. Compounding the challenge, transcription exhibits significant extrinsic and intrinsic noise [19–21]. In particular, the stochastic nature of transcription makes coordinating expression across multiple genetic elements challenging [22–24]. Spatial variation in the nucleus and biochemical dynamics in condensates may contribute to bursting but provide limited parameters for tuning transcriptional noise [25, 26]. Alternatively, mechanical sources of gene regulation offer one potential mechanism by which to understand and harness transcriptional noise to improve the predictable design of gene circuits [27–31].

The mechanical forces of DNA supercoiling powerfully shape transcriptional variance [21, 32]. In the process of transcription, RNA polymerases induce a leading wave of positive DNA supercoiling [33, 34], reshaping the local structure of chromatin [35–38]. At the kilobase scale, chromatin structure correlates with gene regulation [39–41]. In yeast and human cells, transcriptionally-induced supercoiling demarks bounds of gene activity [35, 37]. In particular, transcriptional activity dictates the strength of contact domains such as gene-to-gene interactions, indicating a role for transcription in forming and maintaining interactions at the kilobase scale [40, 42]. Together these data suggest that the process of transcription drives formation of supercoiling-linked, kilobase-scale structures that feedback into transcriptional regulation of gene expression. As supercoiling rapidly diffuses across long distances [43], transcriptional activity at one site may impact the overall activity and dynamics of transcription of proximal genes [44–46]. Understanding how supercoiling induces coupling between neighboring genes provides the opportunity to improve the predictable design of transgenic systems from simple reporters to sophisticated dynamic circuits.

Here we develop a model of transcriptional regulation that integrates DNA supercoiling to examine how the orientation and placement of neighboring genes affects expression. Extending from a model of supercoiling-dependent polymerase motion [44], we developed our model to include the effects of supercoiling on polymerase binding and initiation. Specifically, we model RNA polymerase binding and initiation as a function of DNA supercoiling, such that underwound DNA favors RNA polymerase binding whereas overwound DNA limits binding. To extract experimentally testable predictions, we apply our model to simple two-gene systems that include a constitutive reporter and an inducible gene. Using these two-gene systems, we find that DNA supercoiling strongly influences the profile of gene expression and that influence is defined by syntax—the relative orientation and position of genetic elements—and the enclosing boundary conditions. In addition to regulating the output of simulated genes, supercoiling-dependent feedback tunes the size and frequency of transcriptional bursts. To investigate how these tunable parameters may impact synthetic gene circuits, we applied our model to a canonical gene circuit, a synthetic toggle switch constructed with different syntaxes. We find that circuit syntax affects the stability of states and sets biochemical parameters required for bistability including repressor cooperativity and RNA stability. Finally, we explored how DNA supercoiling might support transcriptional coordination within the native genome of zebrafish (*Danio rerio*) to enable somite segmentation. We find that DNA supercoiling acts as a mechanism for coordinating expression between divergently expressed genes in the segmentation network. Supercoiling-dependent feedback supports tight regulation of these proximal clock genes, providing a molecular mechanism for the precise coordination of gene expression observed during somite formation [47]. Thus, supercoiling-mediated feedback represents a novel, testable regulatory mechanism that can both explain native behaviors and guide synthetic designs.

## 2 Model and methods

Simulating the behavior of native and synthetic circuits under the influence of transcription-induced feedback requires a model that integrates explicitly-modeled RNA polymerase (RNAP) motion and RNA- and protein-mediated feedback mechanisms. Our method combines three modeling levels: an ordinary differential equation system that simulates the continuous progression of polymerases loaded onto DNA, a core stochastic model that models supercoiling-dependent polymerase initiation, and a user-specified stochastic layer that allows for simulation of other modes of transcriptional regulation such as the promoter-repressive and -activating interactions that are often included in synthetic circuits.

### Continuous model of supercoiling-dependent transcription

Here, supercoiling is defined as the amount of excess twist *ϕ* relative to relaxed DNA. Relaxed DNA rotates one full revolution per ≈10 basepairs (1bp ≈0.34 nm); thus its relaxed twist is *ω*_0_ = 1.85 radians/nm. Supercoiling density can generally be defined as a varying function of genomic location *σ*(*z*). However, in a region of constant supercoiling density, we can use the excess DNA twist *ϕ*_1_, *ϕ*_*i*+1_ at the endpoints *z_i_, z*_*i*+1_ to define the supercoiling density as:

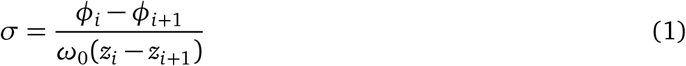

For an polymerase with *ϕ*_1_ > 0 between endpoints with *ϕ*_0_ = *ϕ*_2_ =0, eq. (1) implies that the supercoiling density is *positive* in front of the polymerase and *negative* behind the polymerase (fig.2). Following on the work of Sevier & Levine [44], we assume that on the length scales of synthetic and native circuits of interest (*O*(≈*10kb*)), the supercoiling density is constant in all regions between polymerases and other barriers. Because supercoiling diffusion and plectoneme hopping [43] occur at rates faster than transcription (supercoiling diffusion: 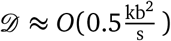 versus transcription rate: 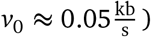)[48], the supercoiling generated by a polymerase will diffuse outward far more rapidly than polymerases can move. We make a pseudo-steady assumption for inter-RNAP supercoiling—assuming that eq. (1) holds between polymerases—over the relatively small (~ 10 kb) genomic distances considered in this work in order to simplify the resulting model.

**Figure 1:**
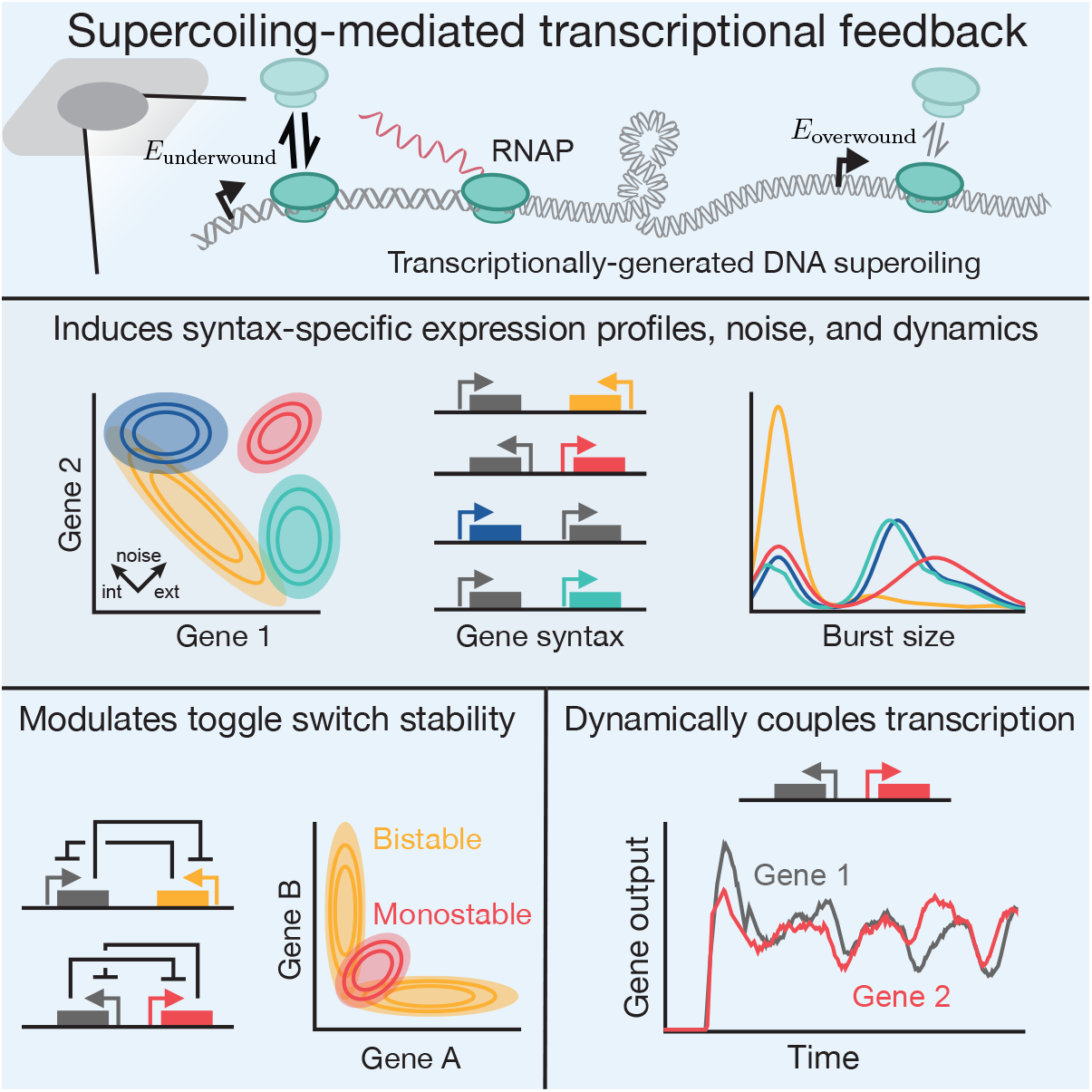
Supercoiling-mediated feedback drives emergent regulatory behaviors. Supercoiling-mediated feedback induces syntax-specific expression profiles and transcriptional variance. Via supercoiling-mediated feedback, circuit syntax modulated toggle switch stability, and dynamically couples transcription through coordinated bursting.

**Figure 2:**
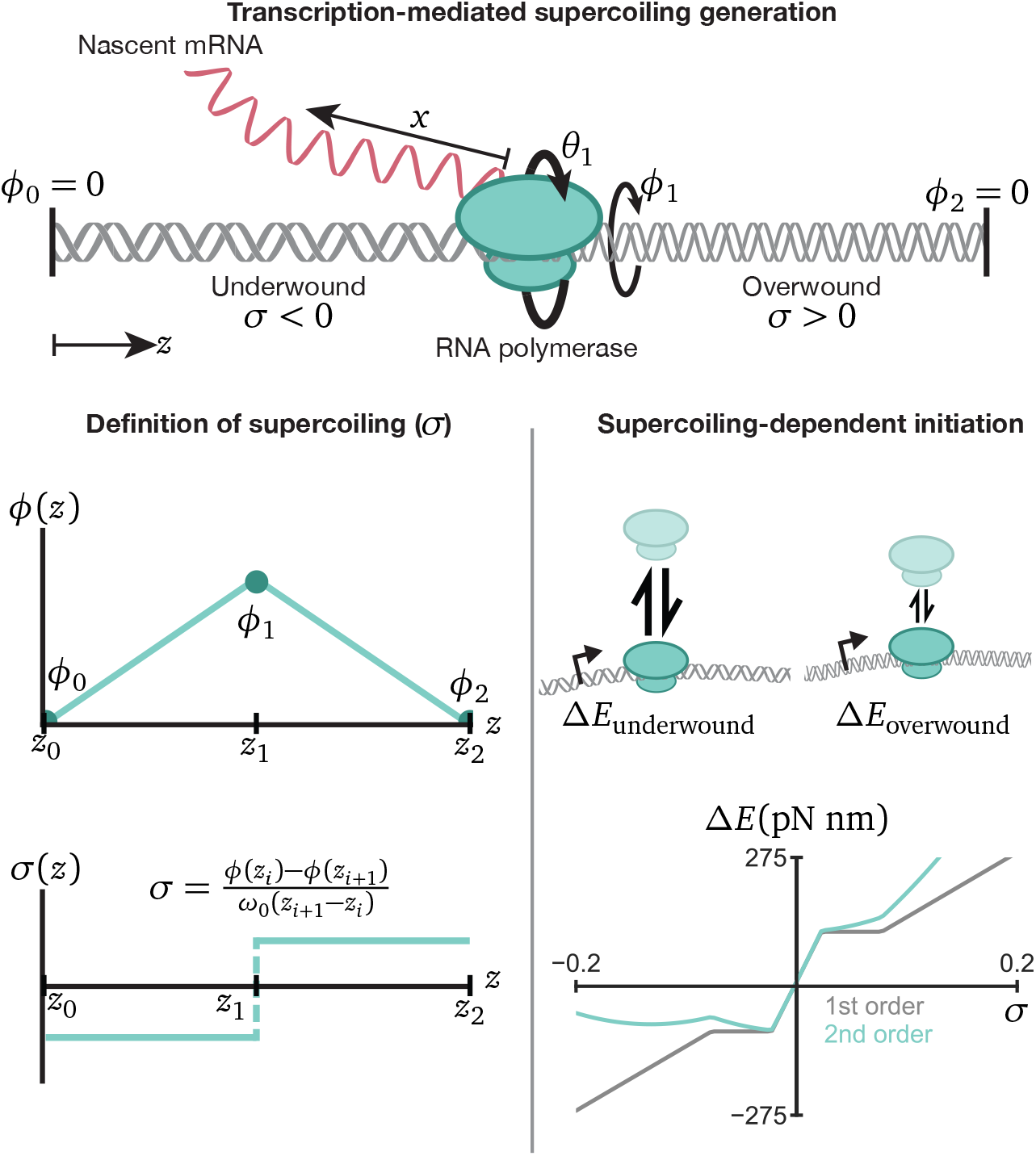
In the limit of fast supercoiling relaxation relative to polymerase motion, the supercoiling density is constant in the region between polymerases and can be calculated from the slope of the linearly-interpolated *ϕ*(*z*) graph. The relaxed DNA twist frequency *ω*_0_ has value *ω*_0_ = 1.85 radians/nm. Four key variables define the location of each polymerase: its linear distance *z* along the genome, the length of the nascent mRNA transcript *x*, the rotation of the polymerase *θ*, and the local DNA excess twist *ϕ*. The tradeoff between RNAP rotation and DNA rotation generates supercoiling upstream and downstream, with the drag generated by the nascent mRNA primarily balancing the torque caused by generated supercoils. Using an energy model responsive to local supercoiling, we can derive supercoiling-dependent initiation terms to model differential polymerase loading rates.

How does transcription both drive the process of supercoiling generation and react to changes in local supercoiling? Under the assumption of supercoiling relaxation, each polymerase is defined by four variables—the one-dimensional genomic location of the polymerase *z_i_*, the length of the nascent RNA *x_i_*, the excess twist at the location of the polymerase *ϕ_i_*, and the rotation angle of the polymerase *θ_i_* (fig.2). Then, two governing equations define the motion of all polymerases [44]. First, we equate linear polymerase motion with the rotational motion required to track the DNA groove:

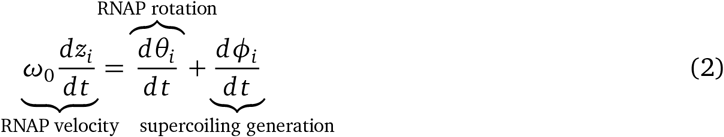

where the change in *θ_i_* representation polymerase rotation and *ϕ_i_* represents local rotation of the DNA. The second equation provides a torque balance between DNA-mediated torques on the left hand side and torque caused by drag acting on the nascent RNA:

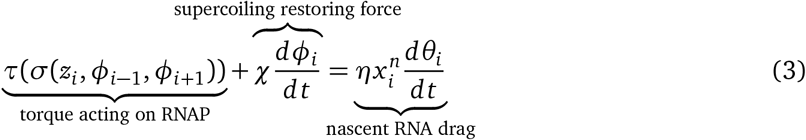

To develop a final system of ordinary differential equations, we still must define the torque response function *τ*(*σ*) and the polymerase velocity function 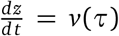. With these two functions, eqs. (2) and (3) can be solved as in Sevier & Levine [44]. Here, we use Marko’s torque-response model to supercoiling which accounts for the thermodynamic behavior of both non-buckled, twisted DNA and buckled, plectonemic DNA (see eqs. (S3) and (S6) in supplemental section C) [49]. The resulting *τ*(*σ*) function exhibits a phase transition, where the torque response is nearly constant at intermediate values of *σ* where the DNA is transitioning from a locally-twisted phase to a plectonemic-phase (fig.S1a).

For the velocity response of a polymerase experiencing a torque *τ_f_* in front and *τ_b_* behind, we model polymerase stalling as:

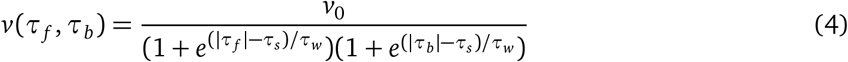

where the stall torque *τ_s_* = 12 pN nm and stall-width *τ_w_* = 3 pN nm define a sigmoidal stall-response curve. As shown in figs.S2c and S2d, our results are only weakly dependent on the specific choice of *τ_s_* and *τ_w_*. Importantly, our selected phenomenological term will stall polymerase motion if *either* the torque upstream or downstream exceeds the stall torque *τ_s_*. Some models choose a stalling equation that only stalls if the difference between the upstream and downstream torque exceeds a stall torque [46]; we chose this form, reasoning that the DNA unwinding and rewinding process opposed, respectively, by upstream and downstream torque could independently stall. When simulated, the difference between these stalling models is small in practice; polymerases at the start or end of the burst encounter both high upstream and downstream torques *and* a high torque difference, whereas polymerases in the middle of a burst experience both low adjacent torques and a low torque difference.

Taking the above equations together, we can simulate the coupled motion of an arbitrary number of polymerases as a single coupled ODE system. We further examine different experimental systems by implementing different boundary conditions that allow us to simulate both plasmid systems and genomically-integrated systems (see supplemental section B.1).

### Modeling supercoiling-dependent initiation

While the described differential equation system can simulate polymerase motion, we need a way to model the addition of polymerases to simulated genes. A simple strategy is to assume a supercoiling-independent initiation rate and use a stochastic simulation method to randomly add polymerases to transcriptional start sites at a certain fixed rate. However, this simple model assumes that polymerases can bind equally well to initiation sites independent of local supercoiling, missing supercoiling-dependent dynamics. In order to include supercoiling in a polymerase initiation model, we relate the basal expression rate to a corresponding base energy term. We can then additively introduce extra energy costs for polymerase binding under different local supercoiling conditions. Under the approximation that the direct energetic cost of locally melting the DNA to fit in the RNAP groove dwarfs the relative change in unwinding energy caused by supercoiling, the majority of the energetic cost comes from inserting supercoiling ahead and behind the inserted polymerase. Under this assumption, the first-order supercoiling energetic correction can be written as:

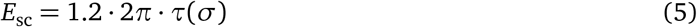

Is this a good approximation? We can estimate the energetic cost of local melting, and find that neglecting local melting leads to a minor change in the resulting energy as seen in fig.S1b. A full derivation of eq. (5) is given in supplemental section C.3.

While this first-order energetic term introduces much-needed behavior to the modeled system—where locally high positive supercoiling decreases the RNAP initiation rate and locally negative supercoiling increases the RNAP initiation rate—at extreme values of *σ*, this energetic term gives aphysical predictions. In particular, under highly negative supercoiling densities, the energetics of polymerase loading becomes increasingly favorable, with loading sometimes occurring more than two orders of magnitude faster when compared to relaxed DNA. To correct for behavior, we add a second-order (quadratic) term of the form that constrains polymerase loading at highly positive or negative local supercoiling:

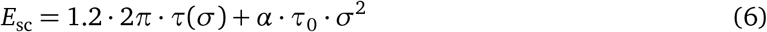

for *τ*_0_, the relevant scale factor in the *τ*(*σ*) equation (eqs. (S3) and (S5) from [49]) and *α*, a positive tunable parameter. As the *τ*(*σ*) equation is linear in *σ* outside of the phase-transition region, this added *σ*^2^ term can be contextualized as an additional term in the Taylor expansion of the physically-realistic *E*_sc_(*σ*) equation. This form of the binding energy enables us to qualitatively match the experimentally observed asymptotic behavior between torque and supercoiling for underwound DNA [50].

For these three models of supercoiling-dependent initiation, we found that the supercoiling-independent initiation model predicted only small changes in reporter output (fig.S3a). Comparing the first- and second-order models, we found that a critical value of *α* existed, *α* ≈ 0.2, above which the second-order model demonstrated emergent non-monotonic behavior (fig.S4). At low values of *α*, the second-order model behaves similarly to the first-order model, so we used *α* = 0.025 for this work. Increasing *α* beyond this chosen value appears to scale down reporter output without qualitatively modifying behavior (fig.S4).

When simulating the ODE model, the rate of stochastic polymerase initiation, *r*_initiation_, varies continuously based on the local supercoiling density *σ* at the transcription start site as:

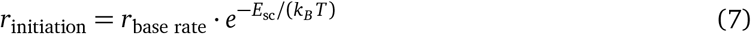

### Modeling additional stochastic, discrete reactions

Many of the native and synthetic systems of interest include mechanisms of gene regulation that rely on other regulatory species. In order to analyze these types of systems using our supercoiling model, we extended our model to simultaneously simulate arbitrary discrete stochastic equations—such as those commonly used in the literature to model protein production, degradation, dimerization, and more. This addition allowed us to model discrete events otherwise not accounted for in the continuous model. Importantly, we simulated the activity of topoisomerases in this way, modeling topoisomerase activity as a stochastic event that removed accumulated supercoiling within intergenic regions.

In addition, we allowed the base initiation rate of genes to vary as an arbitrary function of all species concentrations in the model (*S_i_*), such that eq. (7) becomes:

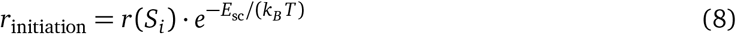

By combining discrete reactions with the ability to dynamically change polymerase initiation rates, we are able to simulate a wide range of phenomena. For example, a cooperative repressive interaction between some repressor protein *R* and a promoter could be modeled using a repressive Hill function:

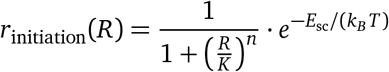

More generally, we can use stochastic formulations of other regulatory mechanisms and test how these mechanisms behave in concert with supercoiling-mediated feedback.

## 3 Results

### Gene syntax and boundary conditions define DNA supercoiling dynamics, expression profiles, and noise

In order to characterize the behavior of supercoiling-mediated feedback, we simulated a series of two-gene systems. These experimentally accessible two-gene systems allow us to test and understand the core design considerations—syntax, relevant boundary conditions, and other experimentally-tunable parameters—within a well-defined and controlled system. Our two-gene systems consist of a reporter gene that is constitutively active and an adjacent, inducible gene placed in either a tandem orientation with the reporter upstream, tandem orientation with the reporter downstream, convergent orientation, or divergent orientation (fig. 3a).

**Figure 3:**
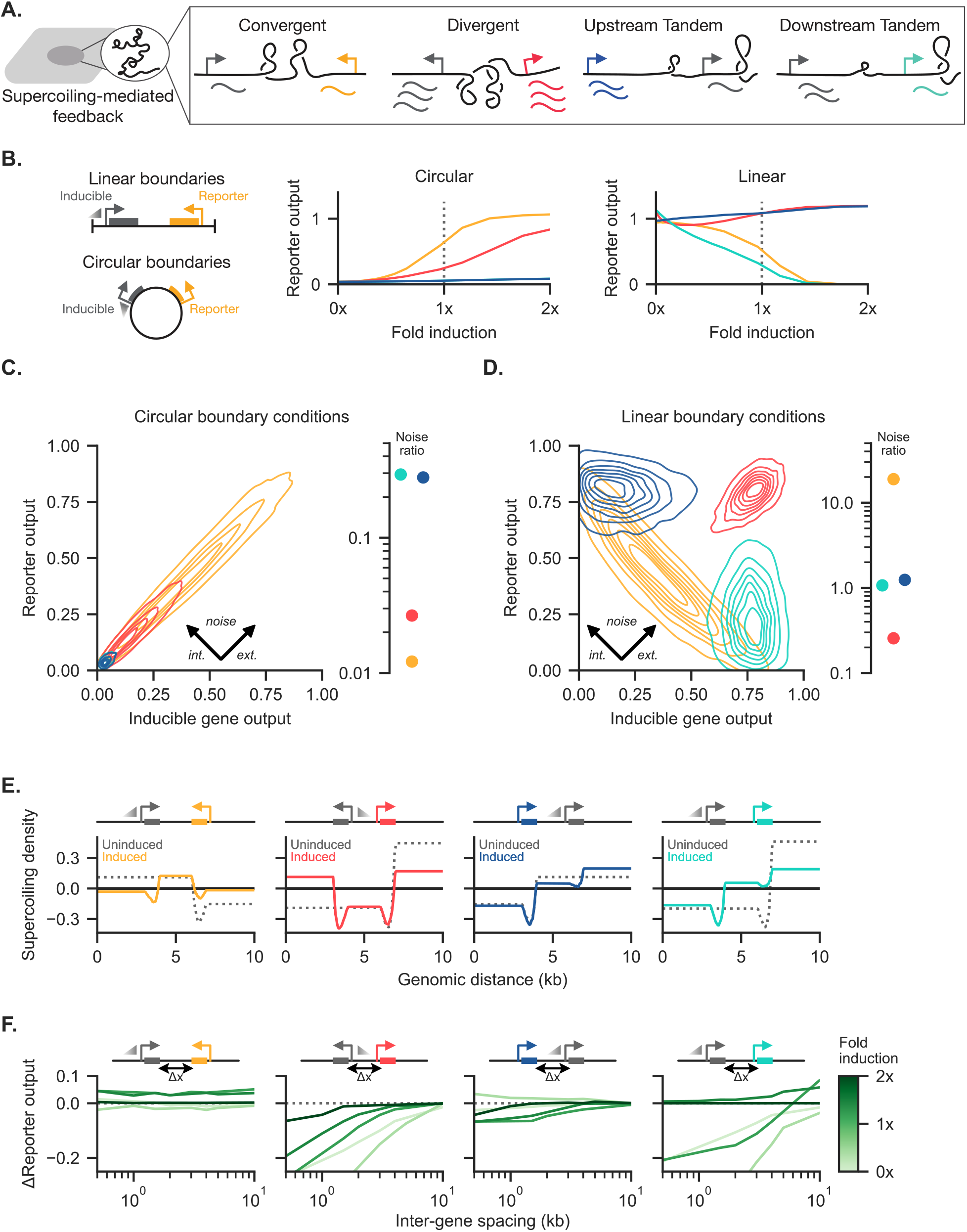
Supercoiling-dependent feedback induces syntax-specific expression profiles. a) Two-gene circuits serve as a testbed for investigating supercoiling-mediated feedback. All four syntaxes include a reporter gene (colored) and an inducible gene (gray). b) Two classes of boundary conditions are simulated. Linear boundary conditions are simulated with adjacent “walls” that prevent supercoiling propagation, whereas circular boundary conditions allow supercoiling generated at one gene to freely affect other genes in either direction around the circle. At right, reporter expression is plotted as a function of the level of induction of the adjacent gene for circular and linear boundary conditions. Reporter output is normalized by dividing mRNA counts by a constant value (250 mRNAs). c) Gene expression distributions for ensembles of circular-boundary-condition simulations are shown, where the adjacent induced gene is equally induced relative to the reporter gene. For each of the four syntaxes, the expression variance can be decomposed into intrinsic and extrinsic noise components; the ratio of intrinsic to extrinsic noise is shown on the right. Reporter output is normalized by dividing mRNA counts by a constant value (340 mRNAs). d) Gene expression distributions and the intrinsic to extrinsic noise ratios are shown for ensembles of simulations with linear boundary conditions. Reporter output is normalized by dividing counts by a constant value (340 mRNAs). e) The supercoiling density across the linear constructs are shown as a function of induction of the inducible gene. Induction (colored line) displays syntax-specific behavior compared to the uninduced case (dashed line). f) Deviations in reporter output relative to the maximum intergene-spacing plotted as a function of intergene-spacing (Δ*x*) for varying levels of induction.

Varying syntax, we examined how boundary conditions affect the expression profiles of the reporter and inducible genes. The type of boundary condition determines how transcriptionally-generated supercoiling propagates to adjacent genes. Experimentally, plasmid DNA and genomically-integrated cassettes allow for interrogation of circular and linear boundary conditions, respectively. Plotting reporter expression as a function of adjacent gene induction, we observed that circular boundary conditions show a monotonic increase in reporter output that scales by syntax (fig. 3b). With circular boundary conditions, the convergent and divergent syntaxes have similar topological layouts, differing only in the relative lengths of the intergenic and inter-promoter regions; as such, the two behave similarly. In contrast, linear boundary conditions show diverse behaviors. The upstream tandem and divergent syntaxes maintain high expression levels while the convergent and downstream tandem syntaxes show decreasing reporter expression with increasing induction of the neighboring gene.

Transcriptional noise substantially contributes to variance of gene expression [24], often confounding circuit designs. Thus, designing circuits to respond to or suppress noise may improve circuit performance. To investigate how syntax impacts expression profiles and noise, we simulated the ensemble behavior for circular and linear two-gene systems at equal transcriptional induction (gray dotted line in fig. 3b). To examine different forms of noise, we decompose the population variance into an *extrinsic* noise component that describes how “all” genes co-vary within a cell and an *intrinsic* noise component that describes the inter-gene variance within a cell. Then, we define the noise ratio as the intrinsic noise divided by the extrinsic noise. For the circular boundary condition simulations, we found that all four ensembles are dominated by extrinsic noise (fig. 3c). In contrast, while the tandem syntaxes with linear boundary conditions show approximately equal intrinsic and extrinsic noise, the linear convergent and divergent populations showed diverging shifts in the variance distribution (fig. 3d). Despite similar levels of extrinsic noise, the divergent syntax minimizes intrinsic variation between the two genes while the convergent syntax maximizes intergene variation. Moving forward, we used the linear set of boundary conditions to analyze system behaviors as we observe the richest set of behaviors under these boundary conditions.

To understand the mechanisms that support syntax-specific expression profiles (fig. 3d), we examined the ensemble supercoiling density of our two-gene systems. Putatively, differences in supercoiling across the two-gene systems give rise to differences in RNAP initiation and thus impact gene expression. To observe supercoiling across the two-gene systems, we averaged the supercoiling density across the simulated ensemble. To examine how induction of the adjacent gene changes supercoiling density, we compared the profiles for when the adjacent gene is uninduced (zero-fold induction) and induced (onefold induction). As expected, the uninduced cases uniformly show that positive supercoiling accumulates upstream of the constitutively-active reporter gene while negative supercoiling accumulates downstream (fig. 3e, for other induction levels, see fig.S5). Upon induction of the adjacent gene, supercoiling accumulates within the intergenic regions in a syntax-specific manner.

Supercoiling *density* determines both supercoiling-dependent initiation and polymerase stalling. Changing the intergene spacing directly tunes the transcriptional activity required to reach a specified supercoiling density and thus reach different expression profiles. To understand the spacing-driven deviations in reporter behavior, we simulated our linear two-gene circuits with different intergene spacings from 500 bp to 10 kb and plotted the output of each circuit, relative to the 10 kb case. Changes in inter-gene spacing mildly influence expression in the tandem and convergent syntaxes (fig. 3f). However, across most induction levels of the adjacent gene, the divergent syntax shows a strong *reduction* in reporter output at small intergenic spacings, potentially demonstrating a tradeoff between the accumulated negative supercoils’ ability to enhance polymerase loading and the ability to stall polymerases due to extreme supercoiling values.

### DNA supercoiling dynamics confer rapid, tunable coupling between adjacent genes

To explore the impact of DNA supercoiling beyond the mean ensemble behavior, we investigated the dynamic behavior of individual simulations upon induction (fig. 4a). To assess how induction impacts dynamics between genes with different syntax, we initialized our two-gene constructs with only the constitutive reporter gene active. After a settling period (2.8 hours), we induced transcription of the adjacent gene. We observe that single-simulation dynamics vary extensively by syntax (fig. 4a). In the convergent syntax, we observe anticorrelated dynamics, with “either-or” promoter activity. In contrast, the divergent syntax supports high levels of expression from both genes. When comparing the two tandem orientations, we observe strong biasing of expression towards the upstream gene with stochastic bursts of expression from the downstream gene. While this upstream dominance does not completely disable the downstream gene, activation of the upstream gene reduces the average expression of the downstream reporter. In addition, we verified that systems initialized with one active gene recapitulate the mean ensemble behavior at steady state (fig.S6), indicating that the systems converge to the same averages independent of initial condition. In other words, our system appears to be ergodic, and we can observe relevant dynamics when initializing the system with only the reporter gene active.

**Figure 4:**
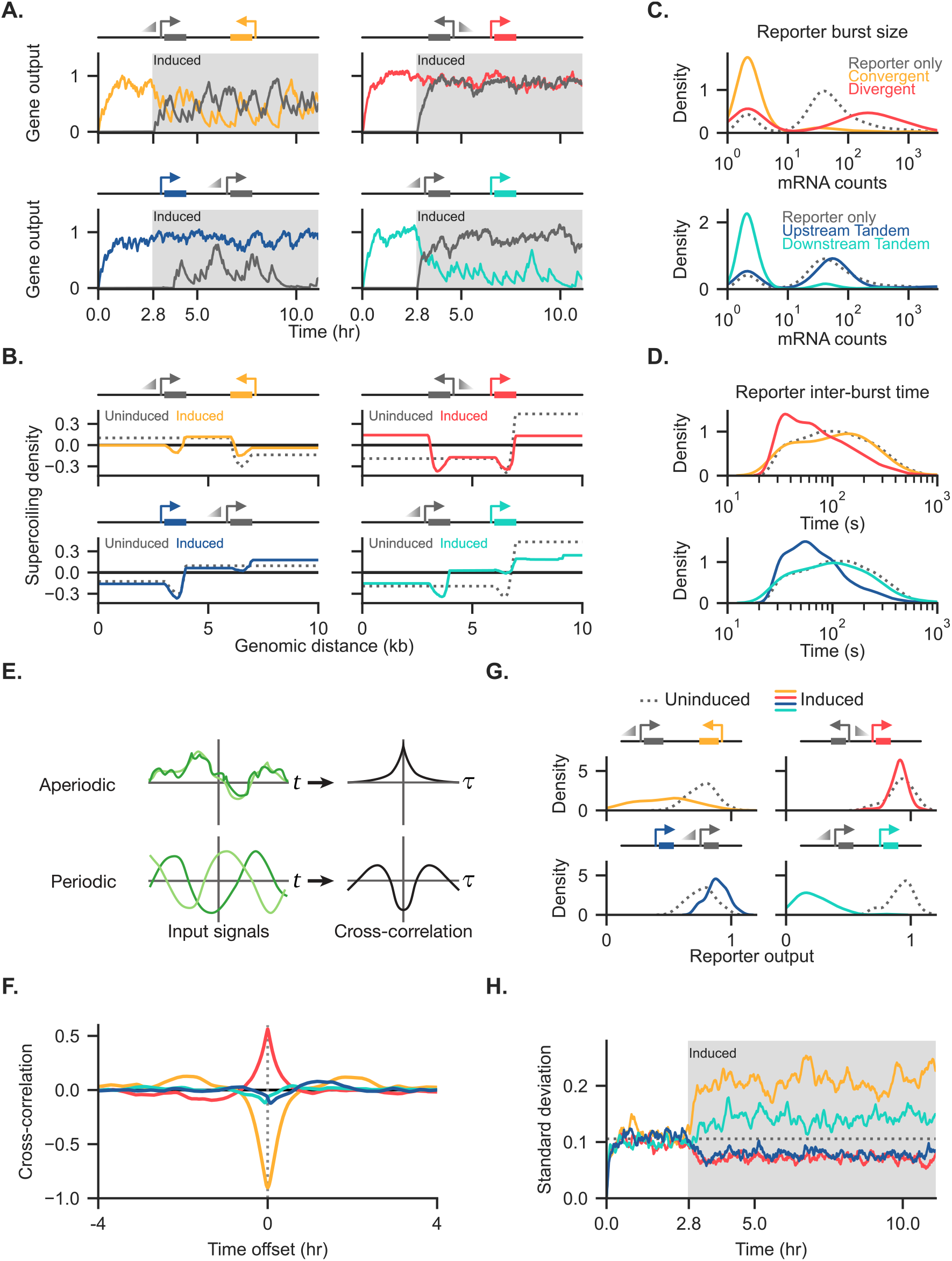
Supercoiling-dependent feedback induces dynamic coupling and altered variance between genes at a single-cell level. a) mRNA counts over time from representative simulations for the four syntaxes. Simulations are initialized with only the reporter gene (colored) active, with the adjacent gene (gray) enabled with equal basal expression after ten thousand seconds (2.8 hours). b) The average ensemble supercoiling density is shown both before and after adjacent gene induction. c) The ensemble distribution of burst size, which is defined as the number of RNA polymerases added per burst, is shown for the different orientations. d) The ensemble distribution of inter-burst time, defined as the time between successive bursts, is shown for the different orientations. e) The cross-correlation of two signals *f* (*t*), *g*(*t*) at a time offset *τ* can be calculated by ‘sliding’ one mean-centered signal relative to the other mean-centered and integrating the product of the resulting signals. f) The cross-correlation between the two genes is shown for the equal-induction case across the four syntaxes. The convergent and divergent syntaxes showed the strongest cross-correlation, with the convergent case showing periodic behavior and the divergent case showing strong correlated expression. g) Distributions of the reporter output before (*dotted*) and after (*solid)* induction of the adjacent gene show changes in both the mean and standard deviation due to adjacent expression. h) Ensemble noise behavior for the four simulated syntaxes is shown by plotting the standard deviation of the reporter gene as a function of time.

When we plot the ensemble-averaged supercoiling density before and after adjacent induction in fig. 4b, we observe similar behavior to fig. 3e, where positive and negative supercoiling accumulate in the intergenic region of the convergent and divergent syntaxes, respectively. Positive supercoiling accumulates downstream in the tandem syntaxes.

Transcription occurs in bursts of activity, and native and synthetic mechanisms can modify burst dynamics [21, 32, 51]. In our model, stochastic polymerase addition leads to transcriptional bursting. In order to understand how circuit syntax contributes to supercoiling-mediated burst dynamics, we examined the distribution of both *burst size*, which we define as the number of polymerases added during a burst, and *inter-burst time*, the amount of time separating two consecutive bursts for the reporter gene (see supplemental section B.4 for more detail). Upon induction of the adjacent gene, we find burst dynamics differ by syntax (fig. 4c). The downstream tandem and convergent syntaxes show a strong decrease in burst size, which we attribute to reduced polymerase loading as a result of increased accumulated positive supercoiling at the reporter promoter region. In contrast, induction of the adjacent gene in the divergent syntax increases the burst size of the reporter gene, putatively due to enhanced loading of polymerases facilitated by accumulated negative supercoiling. Examining the inter-burst time distributions, the upstream tandem and divergent syntaxes show a shift to shorter inter-burst times (fig. 4d). Interestingly, the inter-burst time in the downstream tandem and convergent syntaxes appears unaffected by adjacent induction. Together these observations suggest that syntax provides a parameter for orthogonally tuning burst size and frequency.

To expand our understanding of the emergent supercoiling-dependent dynamics, we examined the temporal correlation of transcription between both genes. To quantify the correlation and extract temporal patterns, we computed the cross-correlation between the gene outputs following induction. The cross-correlation of two signals is itself a function of a time offset; the cross-correlation at some offset time *τ* can be thought of as the Pearson correlation coefficient between the two signals where one has been shifted by *τ* (fig. 4e, see supplemental section B.3 for more detail). In particular, periodic but out-of-phase signals appear as strong negative and positive peaks on a cross-correlation plot. The time offset of peaks corresponds to the phase offset between the signals.

The four syntaxes show starkly different cross-correlation behavior (fig. 4f). The convergent system shows a very large anti-correlation at zero time offset with positive correlation peaks at offsets around ±2 hours. This combination suggests a periodic but out-of-phase behavior between the two genes with a period of around two hours, confirming that our ensemble behaves similarly to the example simulation in fig. 4a. In contrast, the divergent syntax shows a strong positive peak at zero time offset, showing strong, aperiodic but correlated behavior. Tandem syntax showed weak correlation between genes.

We then examined how adjacent induction affects the reporter output distributions. We find that the behavior exemplified in fig. 4a generalizes to the entire ensemble (fig. 4g). The downstream tandem and convergent cases show a reduction in reporter output, the upstream tandem case shows a small increase in reporter output, and the divergent reporter output stays relatively constant. At high adjacent induction, we observe that the upstream tandem and divergent cases show enhanced transcription, with the downstream tandem and divergent cases effectively turning off (fig.S7). We also noted that the width of the distributions change before and after induction, suggesting a change in the noise profile.

As noise impacts the properties of native and synthetic gene networks, we quantified the width of the these distributions by plotting the standard deviation of the ensemble reporter output as a function of time (fig. 4h). Prior to induction of the second gene, all four systems display similar standard deviations. Syntax differences emerge upon induction of the second gene. We found that syntax strongly modulates the noise behavior of the reporter. In particular, the downstream reporter in the tandem syntax and the reporter in the convergent syntax show a strong *increase* in noise levels while the upstream reporter in the tandem syntax and the reporter in the divergent syntax show a small *decrease* in noise levels. These changes in noise can be understood in light of the change in burst dynamics presented in figs. 4c and 4d. The reduction of noise and mean expression value in the downstream-tandem and convergent cases occurs concomitant with a *decrease* in the burst size (fig. 4c). This is expected; if each burst is on average smaller, fluctuations in burst size will have a more dramatic effect on each individual simulation. The decrease of noise in the downstream-tandem and divergent syntaxes is matched by a decrease in the inter-burst time (fig. 4d). Because bursts happen more frequently, the ensemble reporter output is more stable as we approach the always-on limit. Taken as a whole, syntax provides a powerful design parameter for inducing and tuning time-dependent behaviors between genes and shaping output gene distributions.

### Optimizing toggle switch performance and stability through circuit syntax

From oscillators to pulse generators, synthetic circuits aim to precisely coordinate dynamic patterns of gene expression. Yet, emergent dynamics, mediated through DNA supercoiling, may support or impede the performance of dynamic circuits. To examine how supercoiling-mediated feedback influences a dynamic circuit, we applied our model to the classic repressor-mediated toggle switch [8]. Toggle switches are well-characterized both theoretically [8] and experimentally [8, 52, 53]. The behavior of a simple toggle under the additional influence of supercoiling-mediated feedback provides an ideal testbed to understand how syntax influences circuit performance.

A toggle switch can be constructed with two genes that mutually repress each other (fig. 5a). Ideally, such a toggle switch exhibits bistability, generating two stable basins. If modeled with continuous, noise-free equations, a toggle switch will remain within one of the basins based on the initial conditions [8]. However, if we treat the mRNA concentration discretely with a stochastic simulation, the system escapes the stable basin with a certain probability, depending on the size of fluctuations relative to the steadystate values. How does supercoiling-mediated feedback impact this probability of escape? How might supercoiling-mediated feedback interact with mutual transcriptional repression and alter toggle switch behavior? To answer these questions, we simulate the behavior of a two-gene toggle switch with our model for various circuit syntaxes. To establish the conventional dual repressor system used for toggle switches, we abstracted the regulatory interaction of the repressors using a Hill function to define the base promoter initiation rates:

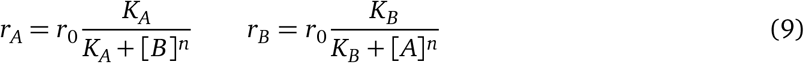

for *r*_0_ = 1/160 s^-1^ and *n* = 2.0. Here, we do not explicitly model protein production. Rather, transcriptional repression directly depends on the mRNA counts of represessors. This parsimonious model allows us to understand the behavior of the system without introducing additional rate constants. We chose the half-max value *K*, the mRNA count at which the promoter activity is half that of *r*_0_, to approximately match the mean steady-state value of either stable state, which ensures that the toggle switch operates in the regime of maximum sensitivity (see supplemental section B.5 for details). In order to compare the behavior of the toggle switches, we initialized toggle switches of different syntaxes within one of the stable basins (gene A) and induced the second gene (gene B) after 2.8 hours (see fig.S9a for example runs). The system then evolves under simultaneous mutual inhibition from expression of the repressors as well as from supercoiling-dependent feedback.

**Figure 5:**
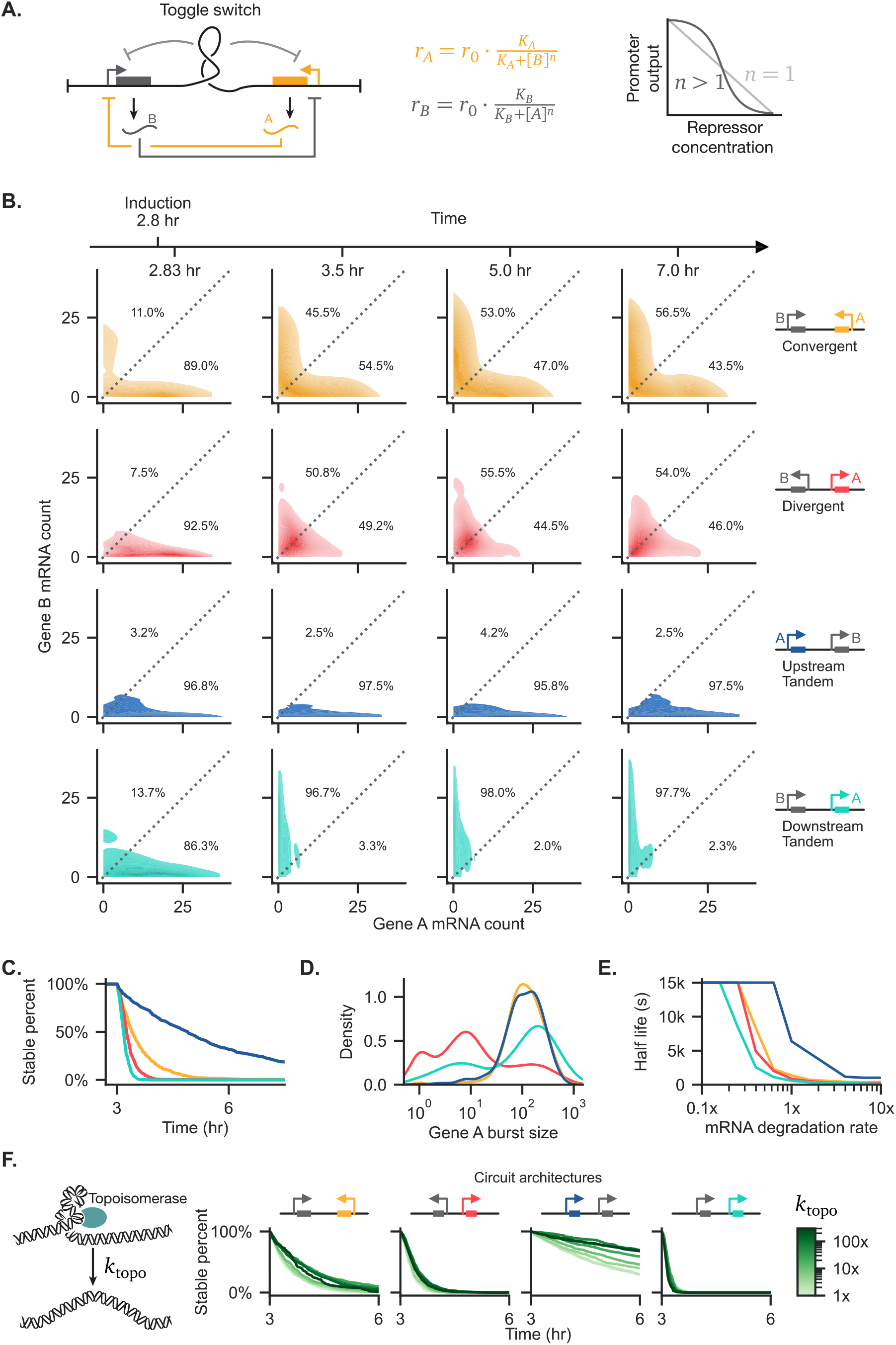
Toggle switches implemented as a mutually-inhibitory pair of genes show syntax-specific stability. a) Schematic of a synthetic toggle switch composed of mutual transcriptional repressors, A and B, which are expressed from a promoter negatively regulated by the opposite gene. Repression follows a Hill function (center), which shows cooperativity based on the value of *n*. Reactions where n is greater than 1 show cooperativity. Simulated toggle switches are regulated both by a mutually-inhibitory interaction at the mRNA level and via supercoiling-dependent phenomena. b) The ensemble mRNA count distributions are shown as a function of syntax at four selected time points. All plots represent simulations where the Hill coefficient has been set to *n* = 2.0. c) The stability, measured as the percentage of simulations in the ensemble that have never escaped the initial starting basin, of the four starting states of the system plotted as a function of time. d) Expression burst size distributions of the initially-active gene A are plotted as a function of circuit syntax. e) The half life at different values of the mRNA degradation rate are shown. As the mRNA degradation rate principally sets the average number of mRNA molecules, high degradation rates lead to systems with low overall mRNA concentration and concordant stochastic instability. f) The stability of the four starting states of the toggle systems are plotted as a function of topoisomerase relaxation rate.

Circuit syntax specifies unique toggle switch dynamics that can be understood by visualizing the distribution of mRNA counts over time (fig. 5b). Initially, nearly all of the simulations in the ensemble lie along the axis corresponding to the initially-active gene, gene A. As time progresses, each ensemble approaches and fluctuates around an equilibrium. The convergent syntax approaches an equilibrium where approximately half of the population distributes into each state. In this syntax, activation of either gene causes positive supercoiling to accumulate in the intergenic region, enhancing negative feedback between genes and thus between states. With divergent syntax, the toggle distributions converge toward monostablity with low differential expression of either gene. Accumulation of negative supercoiling between genes enhances polymerase loading, weakening negative feedback. Finally, the vast majority of the simulations in the tandem syntax remain in or transition to the upstream-active state, demonstrating upstream dominance. We find that these results qualitatively hold for varying values of *n*, the repressor cooperativity coefficient (fig.S10). We find that even in the absence of cooperativity, *n* = 1.0, the convergent syntax shows some level of bistability (fig.S11), indicating that supercoiling-mediated feedback introduces a degree of nonlinearity that can reinforce toggle switch function.

To quantify these distribution results, we computed the *stable fraction* of the ensemble, defined as the fraction of simulations that have never left the initial starting basin at a certain simulation time. The stable fraction monotonically decreases toward zero with time, as simulations that cross into the other stable basin are no longer counted as stable even if they return to the original basin. We observe substantial syntax differences in the dynamics of the stable fraction of the ensemble (fig. 5c). While the tandem orientations represent the extremes of stability, the convergent and divergent syntaxes exhibit intermediate stabilities. As expected, different burst dynamics characterize toggle switch behaviors, varying by syntax. In particular, we observe that the divergent syntax displays *reduced* burst size when compared to the other syntaxes (fig. 5d). We hypothesize that the divergent toggle switch may be governed by a conflicting interaction at the promoter level between supercoiling-mediated feedback and mutual inhibition. Overall, these trends suggest that toggle switch behavior emerges through correlated (or anti-correlated) transcription rather than through differences in burst size and inter-burst time, pointing to potentially orthogonal modes of tuning expression of genes within toggle switches.

For any stochastic system, the steady-state number of molecules influences the stability of the system. As the reservoir of molecules grows larger, the size of fluctuations relative to the total concentration decreases. For toggle switches, we expect that as the number of steady-state mRNA molecules grows, we should approach the theoretical, continuous solution that predicts that no state-switching occurs. To examine this expectation, we modified the simulated mRNA degradation rate, scaling *K* as described in supplemental section B.5, and plotted the resulting half-lives (fig. 5e). As the mRNA degradation rate goes to zero, we increase the reservoir size and observe that the half life for all syntaxes approaches the simulation upper-limit on the half life (fig. 5e). Interestingly, increasing mRNA degradation rates reduces the asymmetry in the half-lives between the tandem upstream and tandem downstream syntaxes. These results suggest that state switching increases as mRNA degradation rate increases as expected.

Finally, topoisomerases regulate DNA topology including DNA supercoiling in the genome, reducing genomic stress. By relaxing accumulated supercoiling across the genome, activity of topoisomerases decreases the average amount of supercoiling. Thus, simulated topoisomerase activity may temporarily relax constraints generated through transcriptionally-induced DNA supercoiling. We examined how the rate of topoisomerase relaxation events affected the stability of the simulated toggle switches. Divergent toggle switches appear insensitive to a large range of topoisomerase rates (fig. 5f). We observe that the most stable systems—the upstream tandem and convergent syntaxes—exhibit topoisomerase-sensitive behavior, showing a minor increase in stability at high topoisomerase rates. Thus, while topoisomerases play an important role across the genome, supercoiling-dependent regulatory behavior may be relatively insensitive to topoisomerase activity.

### DNA supercoiling tightly coordinates expression of proximal segmentation genes

DNA supercoiling provides a mechanism for the precise coordination of colocalized genes. Through colocalization, native circuits may incorporate transcription-linked feedback mechanisms to reduce noise and tune cell-state specific output in tightly-regulated, dynamic processes such as somite formation. In zebrafish, proper somite segmentation requires precise coordination of two clock genes, *her1* and *her7. her1* and *her7* form an inhibitory feedback loop encoded in a divergent syntax (fig.6a). Proper somite formation requires one intact allele of *her1* and *her7*, provided these genes are expressed from the same locus [47]. Mutant zebrafish embryos where *her1* and *her7* are only expressed from separate loci (unpaired) eliminate any supercoiling-mediated coupling while retaining the dimer-mediated inhibitory feedback loop. Zinani *et al*. [47] found that in the unpaired embryos, transcriptional coordination between genes is lost and proper somite segmentation is disrupted (fig.6b). Consequently, physical colocalization represents an important feature supporting transcriptional coordination between genes and proper somitogenesis, which may be mediated by supercoiling-mediated feedback.

**Figure 6:**
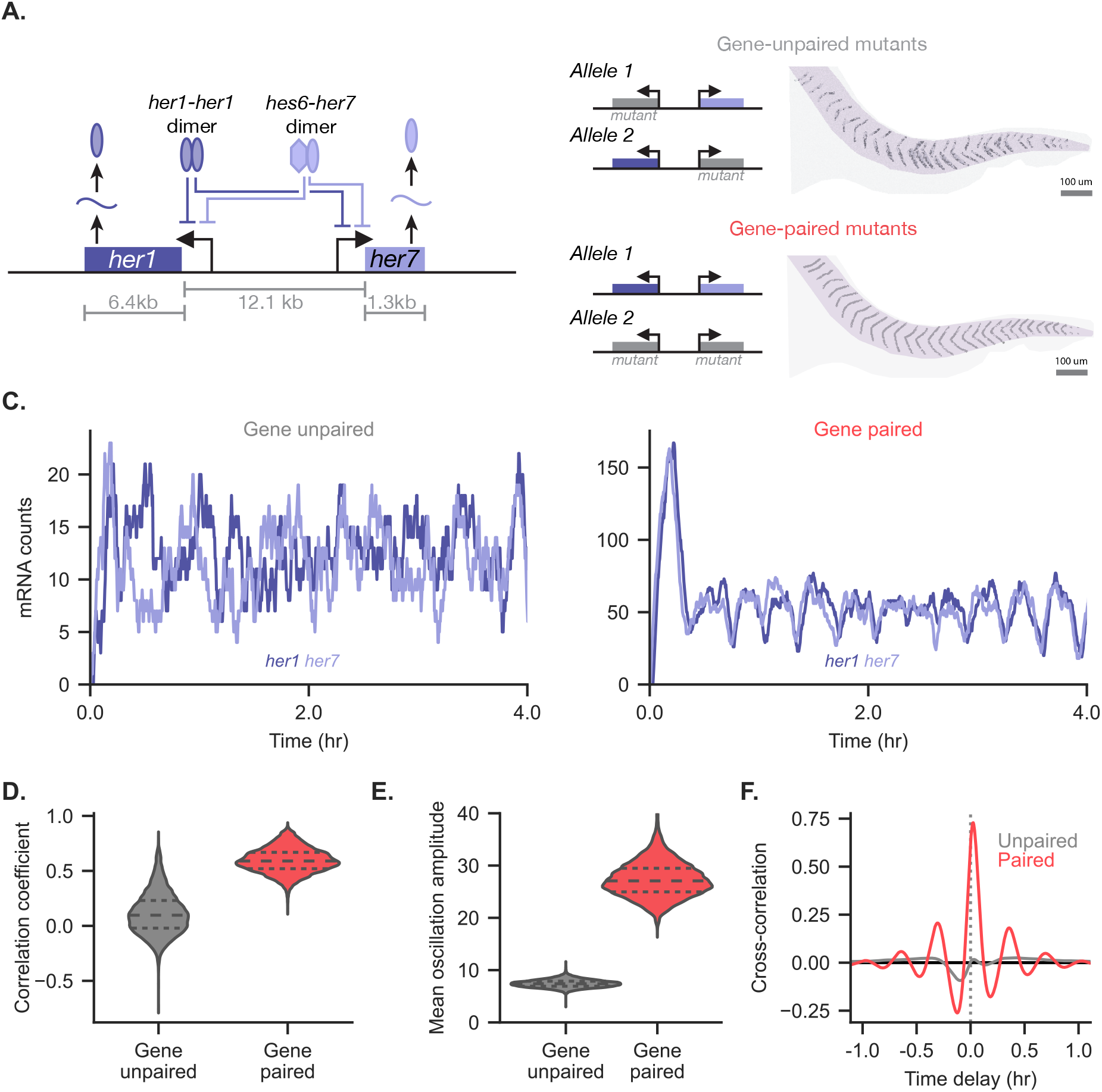
Supercoiling-mediated feedback supports robust transcriptional coordination within the zebrafish segmentation clock circuit.a) Schematic of the mutually-inhibitory *her1-her7* system. Either a *her1-her1* dimer or a *hes6-her7* dimer can bind to either promoter, preventing transcription of the downstream gene.b) Coupling between *her1*, *her7* genes on the same allele supports proper zebrafish somite formation [47]. Disruption of this intra-allele coupling through unpaired mutations in *her1-her7* leads to loss of proper segmentation.c) Two example traces of *her1* and *her7* mRNA levels. The number of *her1* and *her7* mRNAs is shown in example simulation runs for two coupling conditions. The gene unpaired system retains the dimer-driven regulation but is not simulated with biophysical feedback. The gene paired system includes supercoiling-driven biophysical coupling. The initial spike of mRNA in the gene-paired case is a result of starting from an all-zero initial condition; this initial condition was chosen to match literature modeling work [47].d) The correlation between the *her1* and *her7* mRNA counts is shown for the entire ensemble of the coupling conditions.e) The oscillation amplitude over the ensemble is shown for unpaired and paired cases.f) The ensemble cross-correlation between the *her1* and *her7* mRNA counts is shown for the paired and unpaired systems. A large maxima at *τ* = 0 combined with large roughly-symmetric minima can support the strong cyclic behavior observed experimentally in zebrafish.

Based on our above results from two-gene systems, we hypothesized the feedback from DNA supercoiling supports coordination between discrete transcripts expressed from divergent promoters. Using our full computational model, we replicated the previously developed stochastic reaction network. Importantly, *her1* and *her7* are regulated in a binary fashion; the promoters are either completely off when bound by a dimer or expressed at their basal rate when unbound. We simulated two cases: an *unpaired* system where the simulated genes were separated by a large distance (1Mb) to prevent supercoiling interactions, and a *gene paired* system where *her1* and *her7* were spaced at their genomically-active locations. In the paired system, linear boundary conditions were used, with boundaries chosen at the nearest adjacent genes on each side in the zebrafish genome.

Strikingly, the biophysically-coupled case shows strong periodic levels of mRNA expression (fig.6c). In fact, such levels of periodicity are not observed even in the original computational model presented by Zinani *et al*. [47] (fig.S12); we confirmed that this is not simply an artifact of the uniform time resampling performed in order to compare our model behavior to the literature model (fig.S13). The level of periodicity appears sensitive to the second-order polymerase initiation model. Performing simulations below the behavior transition at *α* = 0.02 (see figs.3 and S4), the system displays weak periodicity (fig.S14). Together these results suggest that strong transcriptional coupling via DNA supercoiling requires a specific energy landscape for RNA polymerase binding and initiation in both native and synthetic gene networks.

In order to confirm that these results apply across the ensemble, we examined the ensemble correlation between the counts of *her1* and *her7* mRNA (fig.6d). We found that while the unpaired case shows minimal correlation, the biophysically-coupled case shows strong correlation between the two clock genes. We attribute this strong, periodic correlation to the additional biophysical coupling conferred by the divergent syntax. Notably, we observed that our model predicts an increase in the amplitude of oscillations (fig.6e). *In vivo*, loss of gene pairing reduces oscillation amplitude, leading to improper segmentation [47]. Thus, we propose that supercoiling-mediated feedback offers a mechanism to support robust oscillations in the *her1* and *her7* network for proper somite formation.

Examining the time-dependent nature of the *her1*-*her7* system, we plotted the ensemble crosscorrelation for paired and unpaired genes (fig.6f). Here, we found that in addition to the enhanced positive correlation peak at a time delay of *τ* = 0 seconds, the paired case showed exceptionally strong, nearly-symmetric cross-correlation at positive and negative time offsets. Such cross-correlation is the hallmark of a periodic signal. Thus, both individual examples (fig.6c) and ensemble behavior (fig.6f) show that supercoiling-mediated feedback provides a strong mechanistic driver of intergene coordination in the *her1*-*her7* clock circuit that is inaccessible to solely dimer-mediated regulation.

## 4 Discussion

Transcription induces significant variance in gene expression. At a single-cell level, individual genes are expressed stochastically, with most genes experiencing relatively long periods of quiescence punctuated by bursts of polymerase activity. Phenomenological models of this process based on stochastic probability distributions can provide some insights, but defining the mechanically-regulated physical factors that influence RNA polymerase dynamics will improve existing models of gene regulation and support enhanced design of transgenic systems. Thus, we aimed to integrate a biophysical model of DNA supercoiling into RNA polymerase binding and transcription to examine how DNA supercoiling-mediated feedback influences patterns of gene expression in two-gene systems. Importantly, we sought to use this model to define a set of experimentally-testable predictions as well as lay the groundwork for future modeling across multigenic loci and circuits.

In this work, we developed a model of supercoiling-mediated feedback that captures emergent coupling between neighboring genes to influence expression levels as well as dynamics. This model allowed us to tractably compute polymerase activity at the scale of synthetic circuits (fig.3). Within supercoiling-mediated feedback, we included both supercoiling-dependent polymerase motion terms and supercoiling-dependent polymerase initiation terms. Importantly, this computational framework lays the groundwork for understanding how DNA supercoiling functions as a regulatory mechanism that can be integrated with canonical biochemical models of gene regulation. Using this model, we extracted insights into how mechanical and biochemical regulation combine to generate diverse profiles of expression and support or impede the performance of gene networks.

We find that induction of neighboring genes significantly influences the transcriptional activity of both genes (fig.3). Syntax-specific differences in DNA supercoiling dynamics, expression profiles, and noise emerge due to physical coupling. By computing ensemble supercoiling density and burst dynamics, we observe that changes in burst size and inter-burst time support syntax-specific expression profiles (figs. 4c and 4d). Syntax-dependent biophysical coupling also dramatically affects a range of system behaviors. In particular, supercoiling-mediated feedback is capable of orthogonally modifying adjacent gene expression and burst dynamics, but is itself dependent on inter-gene spacing, mRNA degradation rate, and other variables tunable in an experimental setting. Generally, accumulated negative supercoiling leads to correlated bursting, which occurs concomitant to a decrease in intrinsic noise. In contrast, accumulated positive supercoiling can lead to anti-correlated bursting, which instead enhances intrinsic noise. While the tandem syntax does not lead to large supercoiling accumulation in the intergenic region, we observe upstream dominance, where the upstream gene is more highly expressed than the downstream gene.

Our prediction of burst dynamics complements theoretical and experimental investigations of cooperative interactions of RNA polymerases arising from the beneficial cancellation of positive and negative supercoiling generated by adjacent polymerases [29, 45]. While we do not directly examine transcription elongation rates, we similarly predict syntax-specific differences in expression dynamics, but observe distinct syntax-specific behaviors in our model [45, 46]. We also find that intergenic distance only weakly affects supercoiling feedback [46]. In alignment with experimental work, we find that positive supercoiling accumulates in the intergenic region of convergently-oriented native genes [38]. When combined with sequencing methods that precisely measure nascent mRNA transcription [54], these methods may provide a window into experimental systems in order to test theoretical predictions of our work and others.

The fast timescale of supercoiling-mediated feedback offers access to a uniquely tunable and orthogonal form of gene regulation. In contrast to regulatory mechanisms dependent on relatively long timescales, such as mRNA- and protein-mediated systems, supercoiling-mediated feedback occurs at the timescale of seconds. Polymerases can stall and unstall each other within seconds, while local polymerase loading rates can vary over the course of minutes. By combining the fast dynamic feedback with slower classic feedback mechanisms, circuit regulation can be selectively stabilized or destabilized. We found that specification of syntax within a simple two-gene toggle switch generated diverse behaviors including: a reasonably stable switch, a hypersensitive toggle with hysteresis, and an asymmetric system that preferentially decays towards a single target state. As a rapid mechanism for coordinating transcriptional dynamics, supercoiling-dependent feedback may support intergenic coordination in native systems. Examining the zebrafish segmentation clock, we find that addition of supercoiling-mediated feedback recapitulates the synchronized, periodic expression of the clock genes, *her1-her7* (fig.6).

In deriving our model, we made several simplifying assumptions. Our derivation of the energy function for supercoiling-dependent polymerase initiation adds relevant molecular detail to our model. The second-order correction term reflects the asymptotic relationship observed in *in vitro* assays between torque and supercoiling density for underwound DNA [50]. Inclusion of this correction supports the periodic behavior of the native *her1-her7* clock circuit, suggesting this term may accurately capture regulation *in vivo*. For simplicity, we model the dynamic processes of RNAP binding, initiation, and pause release as a single reaction. In real biological systems, each of these processes may vary across the genome by sequence and by the presence of DNA-binding proteins. Formation of supercoiled structures shows sequence bias *in vitro* which may impact *in vivo* structures and gene regulation [55]. Additionally, our model excludes nucleosome occupancy, polymerase collision and premature termination, and the impact of 3D structures and loop domains formed by protein complexes such as CCCTC-binding factor (CTCF) and other structural maintenance of chromosomes (SMC) proteins. Nucleosomes act as a “store” of negative supercoiling and are displaced by polymerase motion; genomic regions differing in average nucleosome occupancy thus would have a different basal supercoiling density that is ignored in this work. More broadly, we assume that regions of simulated chromatin are uniformly accessible and have equal torque responses during polymerase elongation; these assumptions may fail at the boundaries of chromatin domains. Finally, we also neglect the speed of supercoiling diffusion. While this may be a negligible effect at the scale considered in this work, supercoiling diffusion remains slow at the scale of hundreds of kilobases to megabases.

Our model integrates supercoiling-mediated biophysical feedback with classic gene regulation motifs that are well-studied in native and synthetic contexts. This unified framework brings us closer to an understanding of how supercoiling contributes to transcriptional regulation. We offer testable predictions about the performance of genetic circuits. The predicted changes in reporter output, supercoiling density, and burst dynamics observed in figs.3 and 4 are experimentally accessible with modern sequencing and single-cell imaging technology [38, 54, 56]. Experimental verification of our theoretical results would aid in constructing a mechanistic understanding of how transcription-induced supercoiling couples expression. Harnessing these insights will enable gene regulation at the level of transcription, providing a robust method to control expression dynamics, levels, and noise.

## Code availability

Full simulation and complete data analysis and figure-generating code is available at https://github.com/GallowayLabMIT/tangles_model.

## Acknowledgements

The authors acknowledge the MIT SuperCloud and Lincoln Laboratory Supercomputing Center [57] for providing HPC resources that have contributed to the research results reported within this paper. We also thank Adam Beitz, Ross Jones, Sneha Kabaria, Brittany Lende, Kasey Love, Emma Peterman, and Nathan Wang for helpful suggestions throughout and for their effort in editing this manuscript.

## Funding

Research reported in this manuscript was supported by National Institute of General Medical Sciences of the National Institutes of Health under award number R35-GM143033.

## Supplemental information

### A Supplemental figures

**Figure S1:**
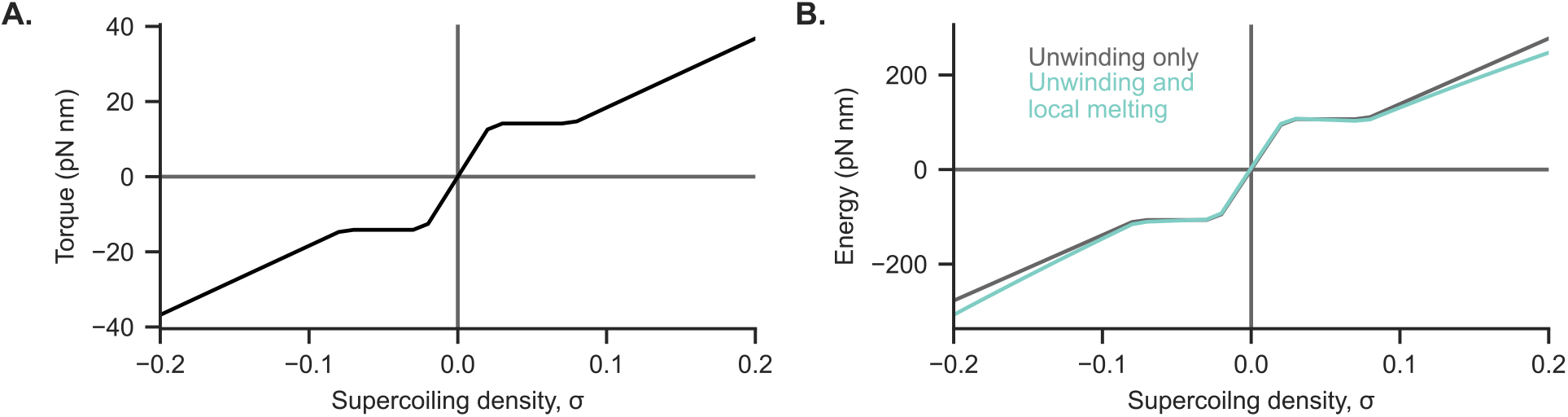
a) The torque response predicted by Marko’s model (eq. (S3)) exhibits phase behavior, with a transition occurring between a over- and under-twisted phase to a largely plectonemic phase. During the transition, the torque response is constant.b) For the supercoiling-dependent initiation model, the energy that it takes to introduce supercoiling upstream and downstream by unwinding the DNA dominates the energy expression. Including a term representing the energy of local melting of the double helix at the location of the polymerase only gives a minor correction.

**Figure S2:**
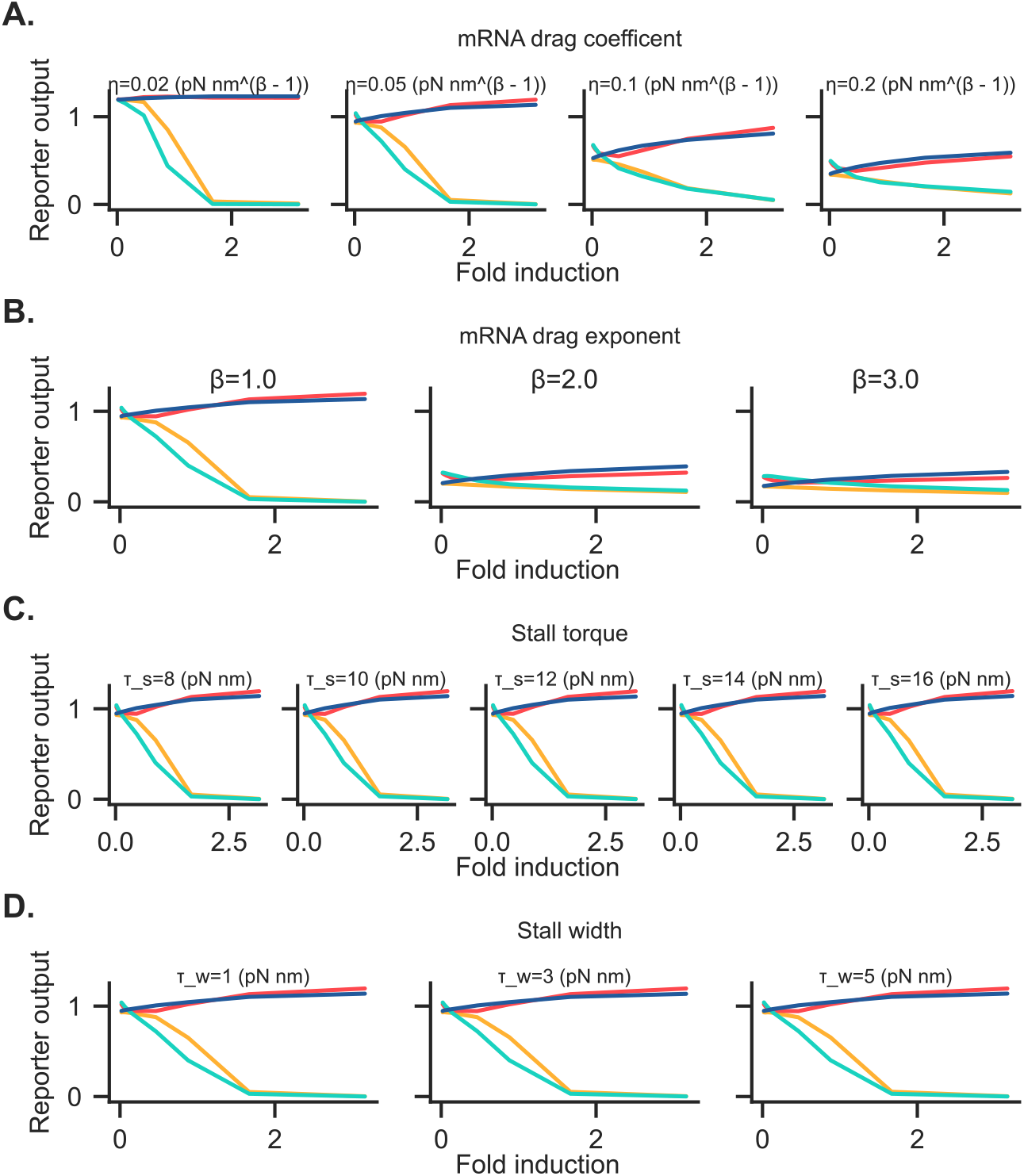
Reporter output of two-gene circuits with linear boundary conditions (as in fig.3) demonstrate similar qualitative behavior when underlying model parameters are changed.a) Varying the nascent mRNA drag coefficient from the chosen value *η* = 0.05 rescales but does not dramatically change reporter output behavior. Larger drag coefficients increases generated supercoiling by introducing additional torque into the system.b) Varying the exponent on the nascent mRNA length in the drag coefficient from the choice of linear drag *β* = 1.0 also rescales but does not dramatically change reporter output behavior. Choices of *β* > 1.0 tends to increase generated supercoiling due to longer nascent mRNAs having an increased drag profile.c) Varying the RNAP stall torque from the chosen *τ_s_* = 12 pN nm has a minor effect on reporter output.d) Varying the width of the stalling function from the chosen *τ_w_* = 3 pN nm has no noticeable effect on reporter output.

**Figure S3:**
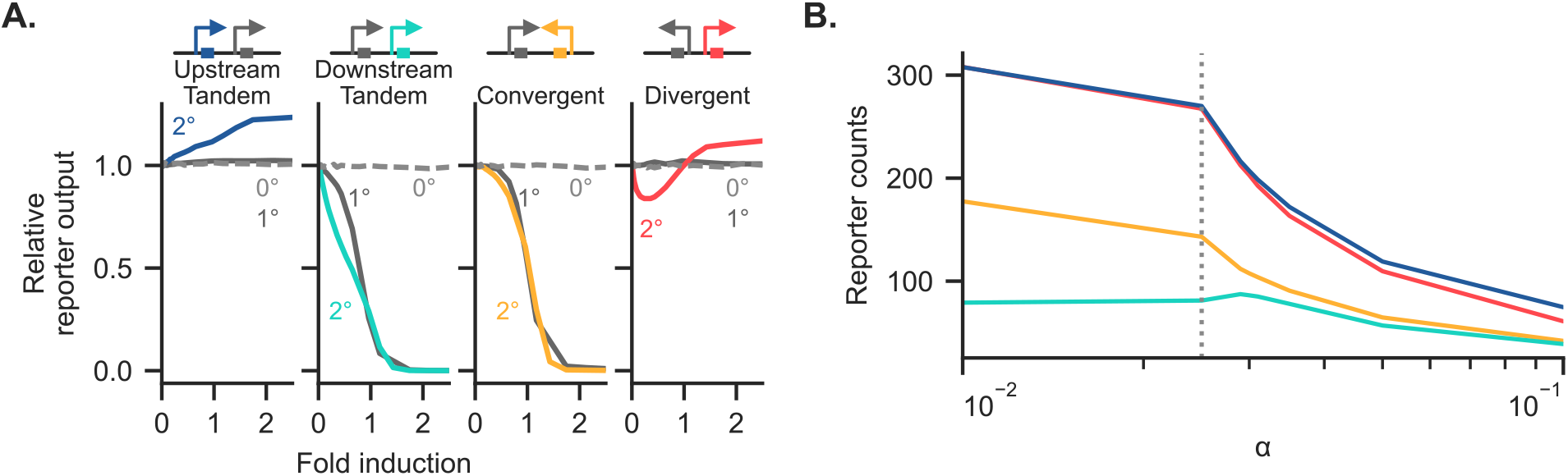
a) Reporter output is shown, normalized to the zero-fold induction case for the four tested syntaxes for the three different polymerase initation models.b) For the second-order model, we chose the value of *α* that defined the transition between regiemes.

**Figure S4:**
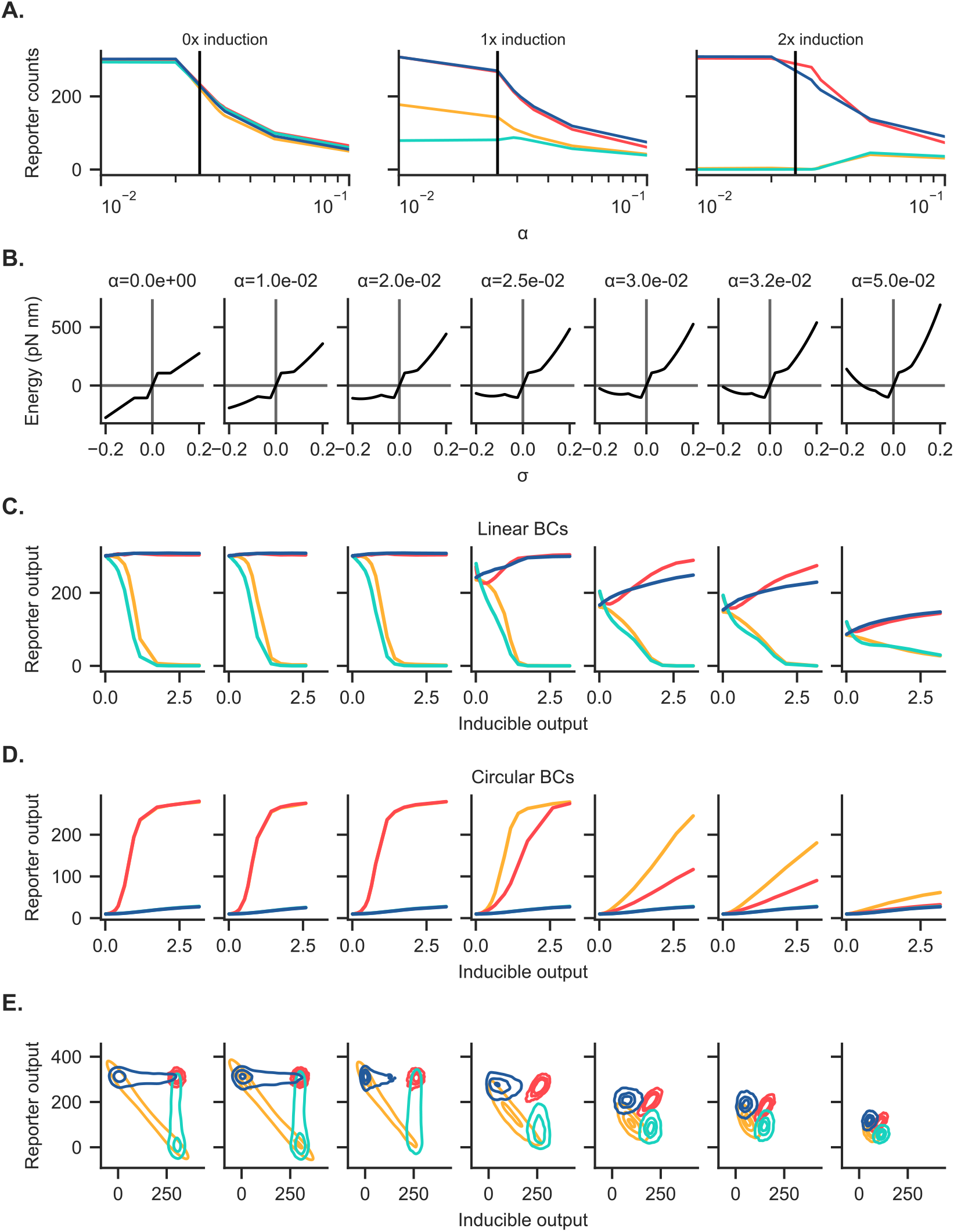
As the value of the quadratic energy correction coefficient *α* varies, two regimes of supercoiling-dependent behavior emerge. For *α* ≤0.02, supercoiling-dependent initiation appears to dominate, leading to both the maintenance of reporter output in the linear divergent case and near-equality of the convergent and divergent cases with circular boundary conditions. For *α* ≥ 0.025, emergent non-monotonic behavior appears, leading to complex behaviors with linear boundary conditions and a separation between the response of the convergent and divergent syntaxes with circular boundary conditions.a) The additional polymerase binding energy as a function of supercoiling density is plotted for various values of *α*.b) c) Reporter behavior of the four syntaxes under linear and circular boundary conditions are shown, in comparison to fig. 3b.d) The overall distribution of the four syntaxes as a function of alpha is shown, in comparison to fig. 3d.

**Figure S5:**
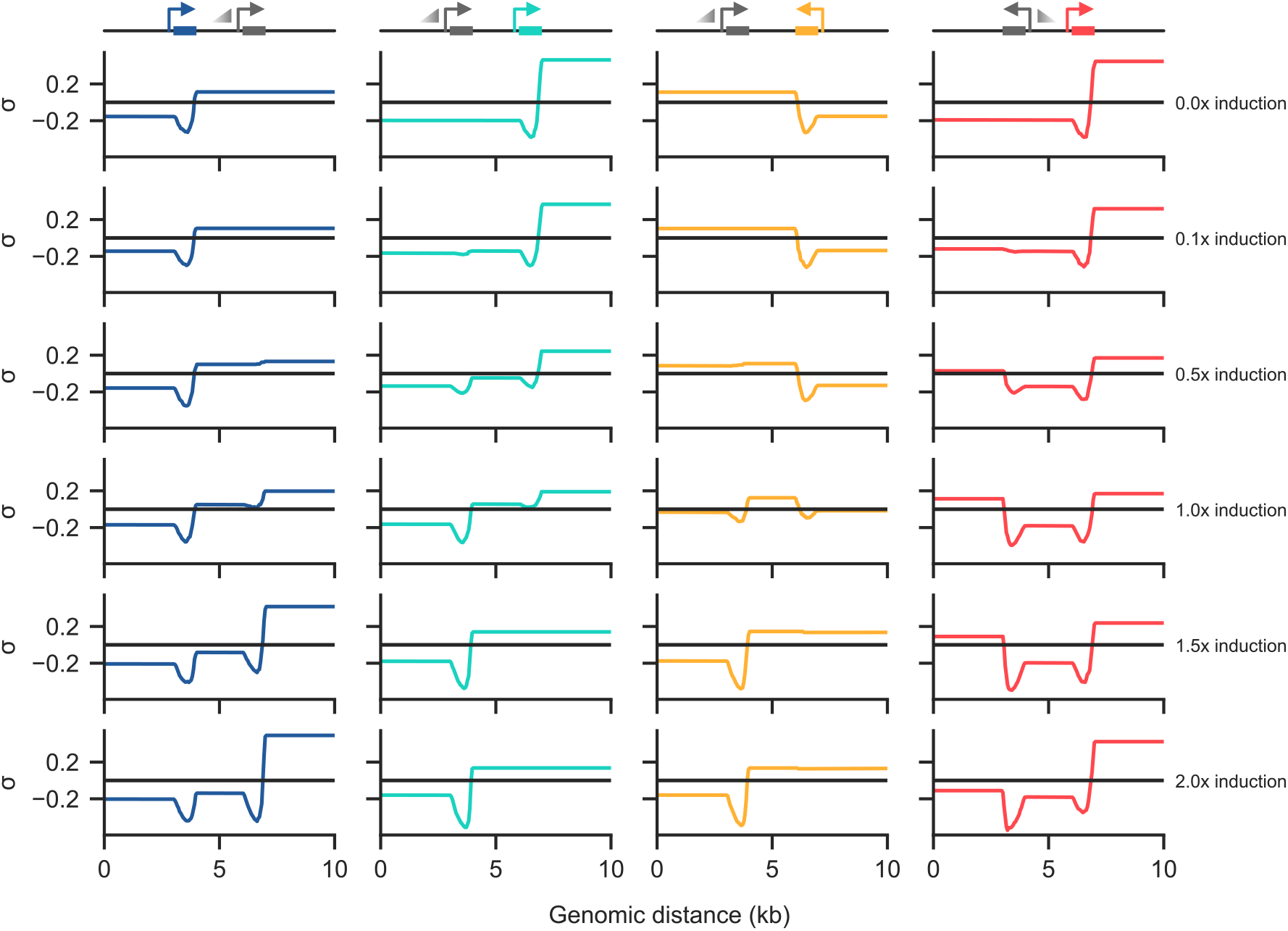
Supercoiling density profiles are plotted for a wide range of adjacent gene induction levels across the four syntaxes considered. At low adjacent induction of the adjacent gene, positive and negative supercoiling mainly accumulates upstream and downstream, respectively, of the reporter gene. At 1-fold (equal base rates) induction, the supercoiling distributions are nearly symmetric in the convergent and divergent syntaxes. At high adjacent induction, the adjacent gene expression dominates the generated supercoiling profiles.

**Figure S6:**
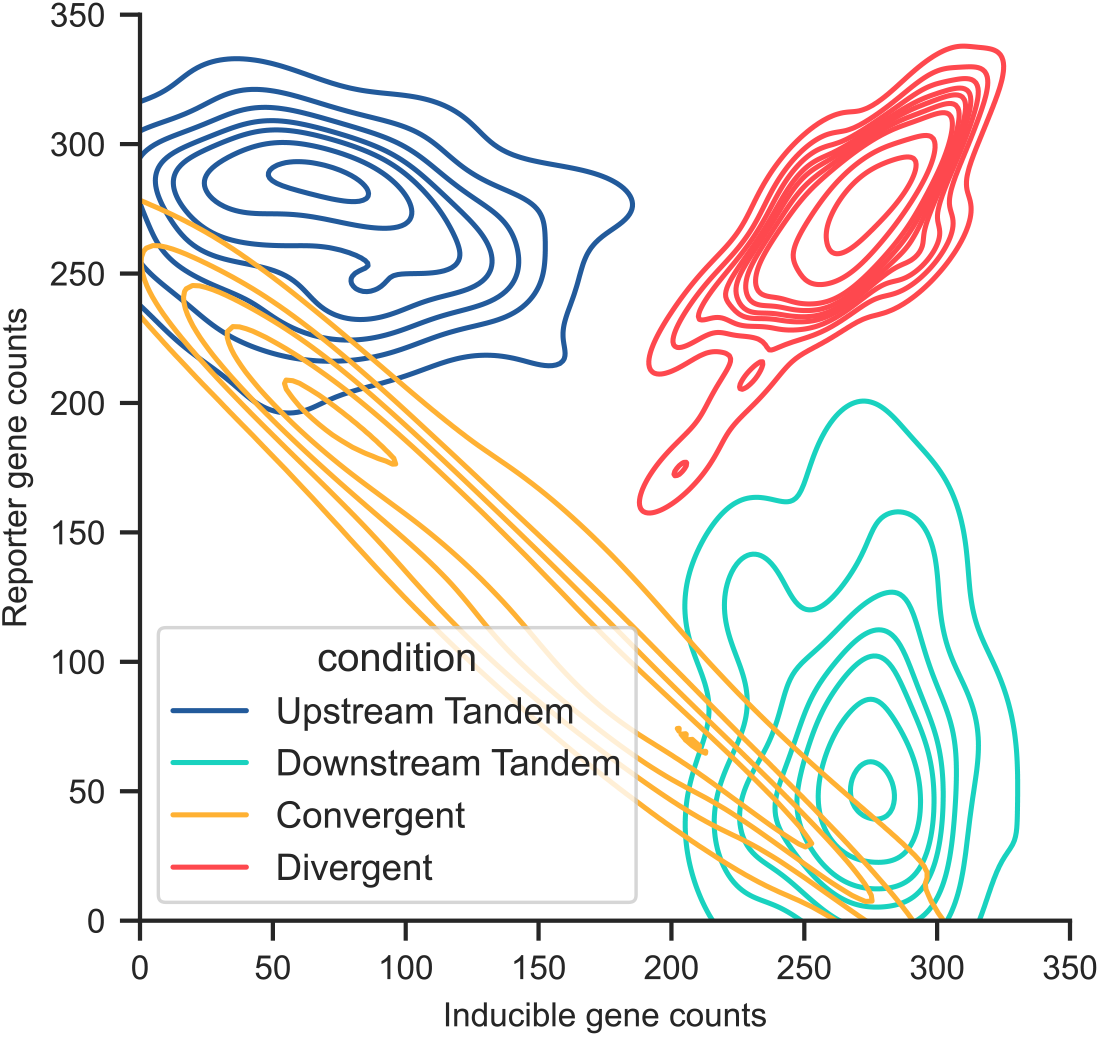
The distribution of inducible and reporter mRNA counts are shown for the simulation ensembles (100 ensembles per condition) presented in fig.4. The different initial starting condition does not affect the final distribution state when compared to fig. 3d, indicating that the final state reached is independent of initial state.

**Figure S7:**
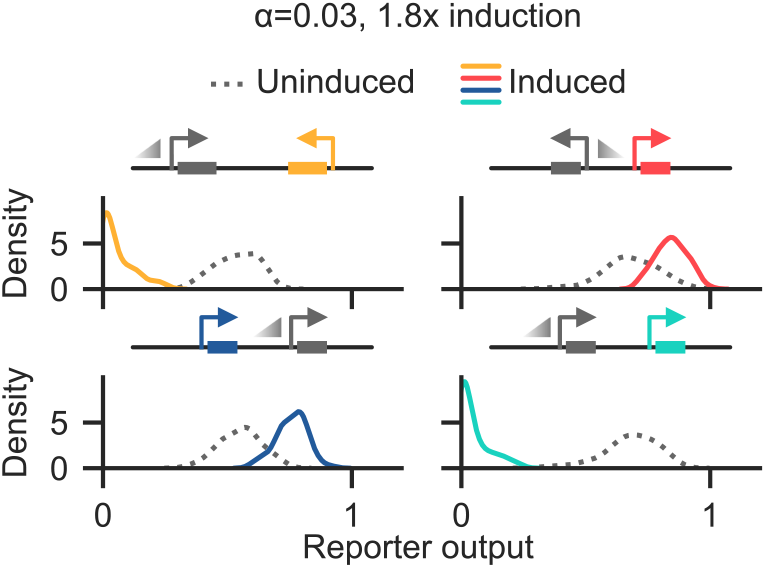
The ensemble distribution of the four linear circuit syntaxes is shown for *α* = 0.03 and high induction of the adjacent gene (1.8 fold).

**Figure S8:**
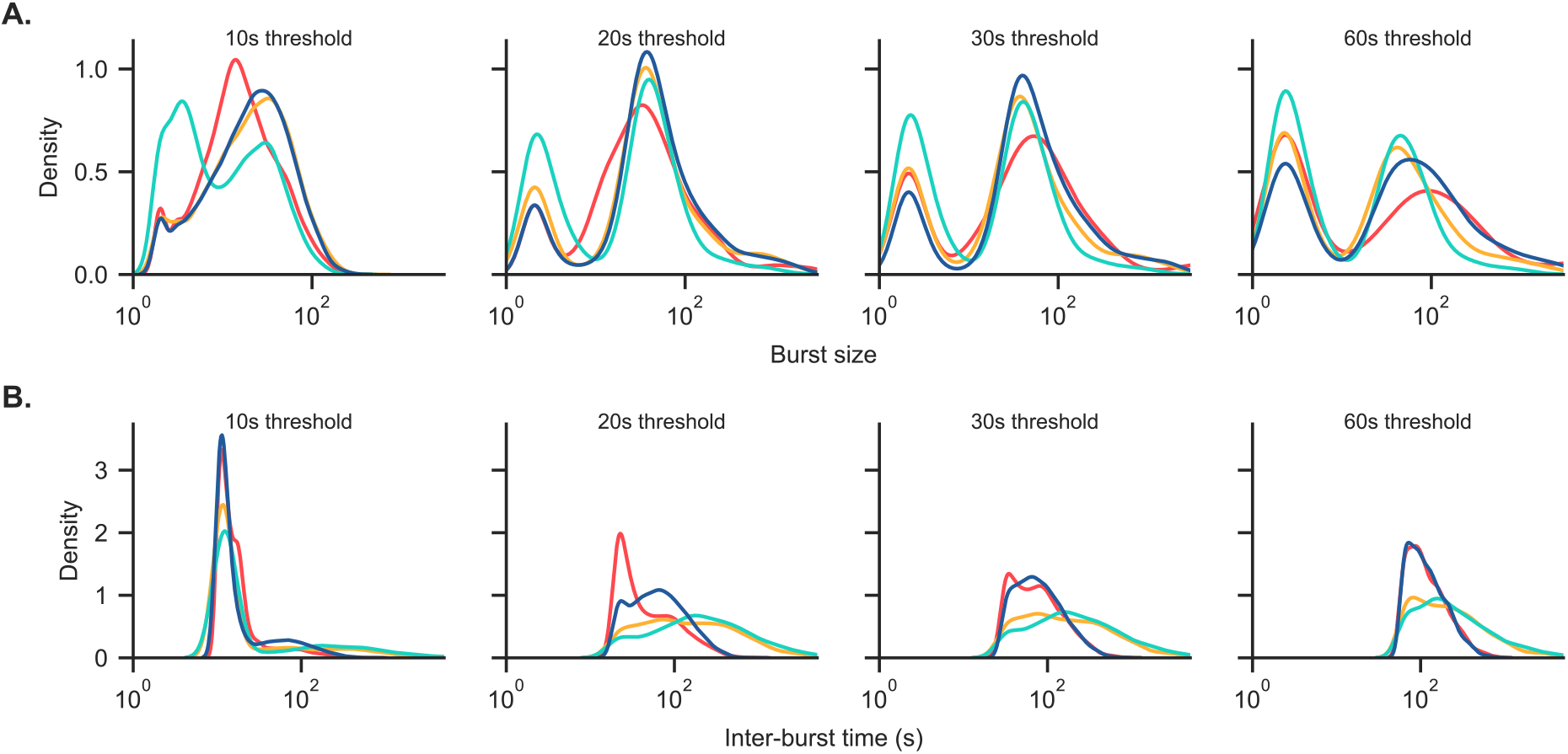
Burst size and inter-burst time distributions are shown at equal-induction as a function of threshold time. At the lowest threshold time, the burst size and inter-burst time distributions qualitatively differ, with only a small percentage of bursts having an inter-burst time larger than 20 seconds. This lack of larger inter-burst times suggests that a burst threshold of at least 20 seconds gives useful predictions.a) Burst size distributions are shown as a function of circuit syntax and burst threshold time.b) Inter-burst time distributions are shown as a function of circuit syntax and inter-burst times.

**Figure S9:**
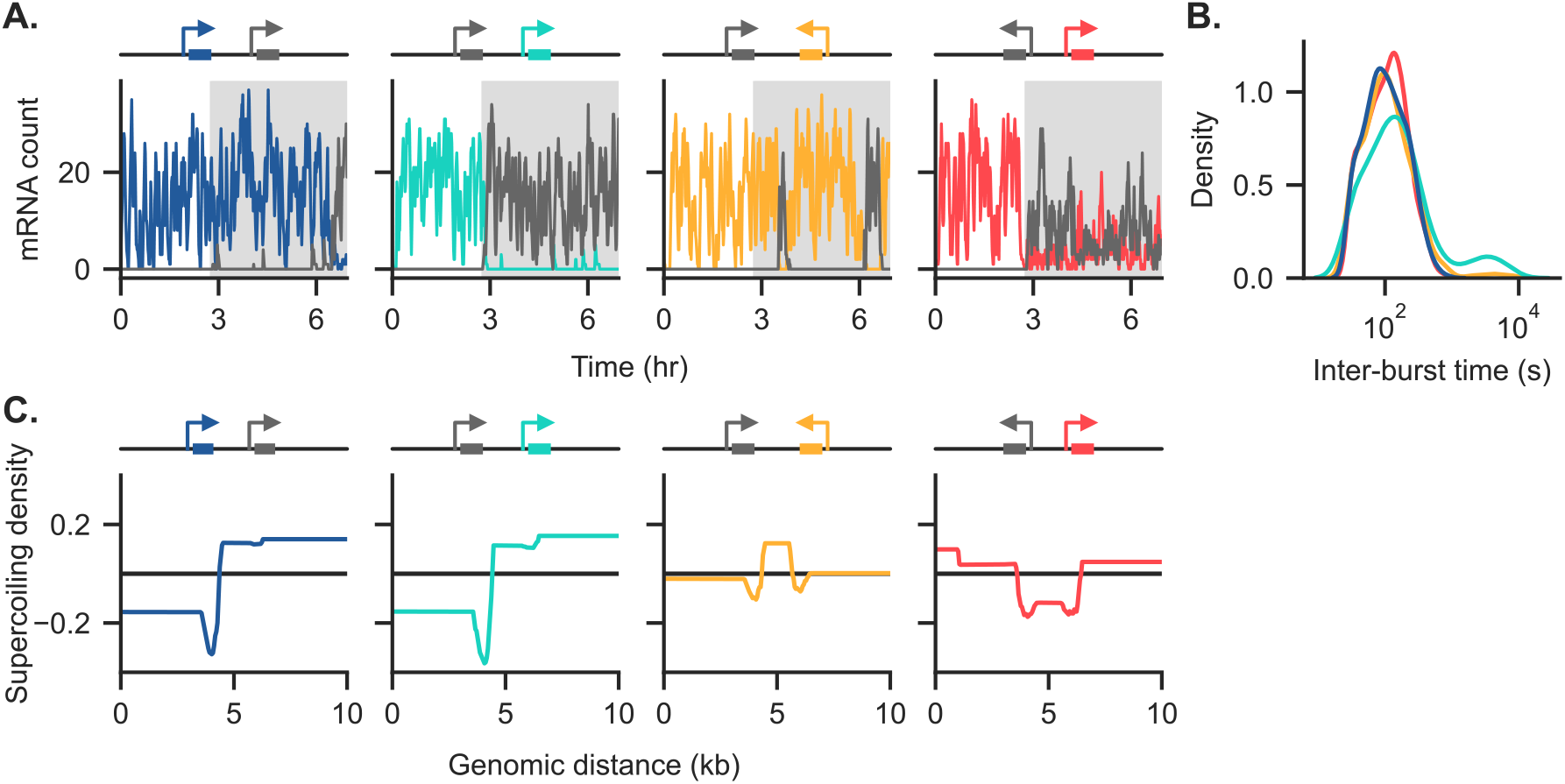
Additional visualizations of the toggle switch circuits presented in fig.5 are shown.a) Four example simulation runs for the toggle switches placed in the four orientations.b) The distribution of inter-burst times for the four different toggle switch architectures.c) The mean ensemble supercoiling density is plotted for the four different circuit syntaxes.

**Figure S10:**
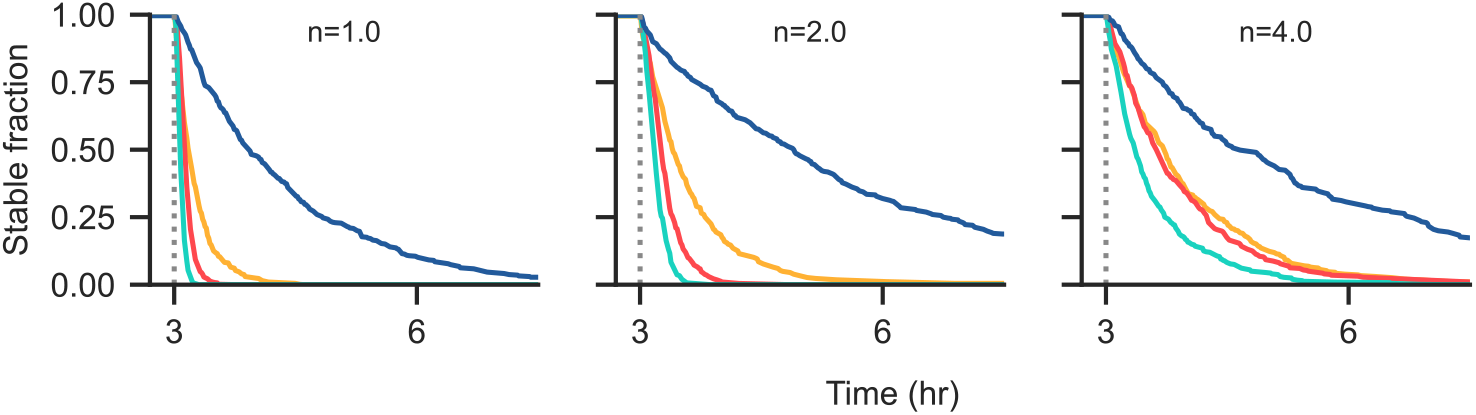
The basin stability for the four toggle-switch architectures is shown explicitly as a function of hill coefficient, *n*.

**Figure S11:**
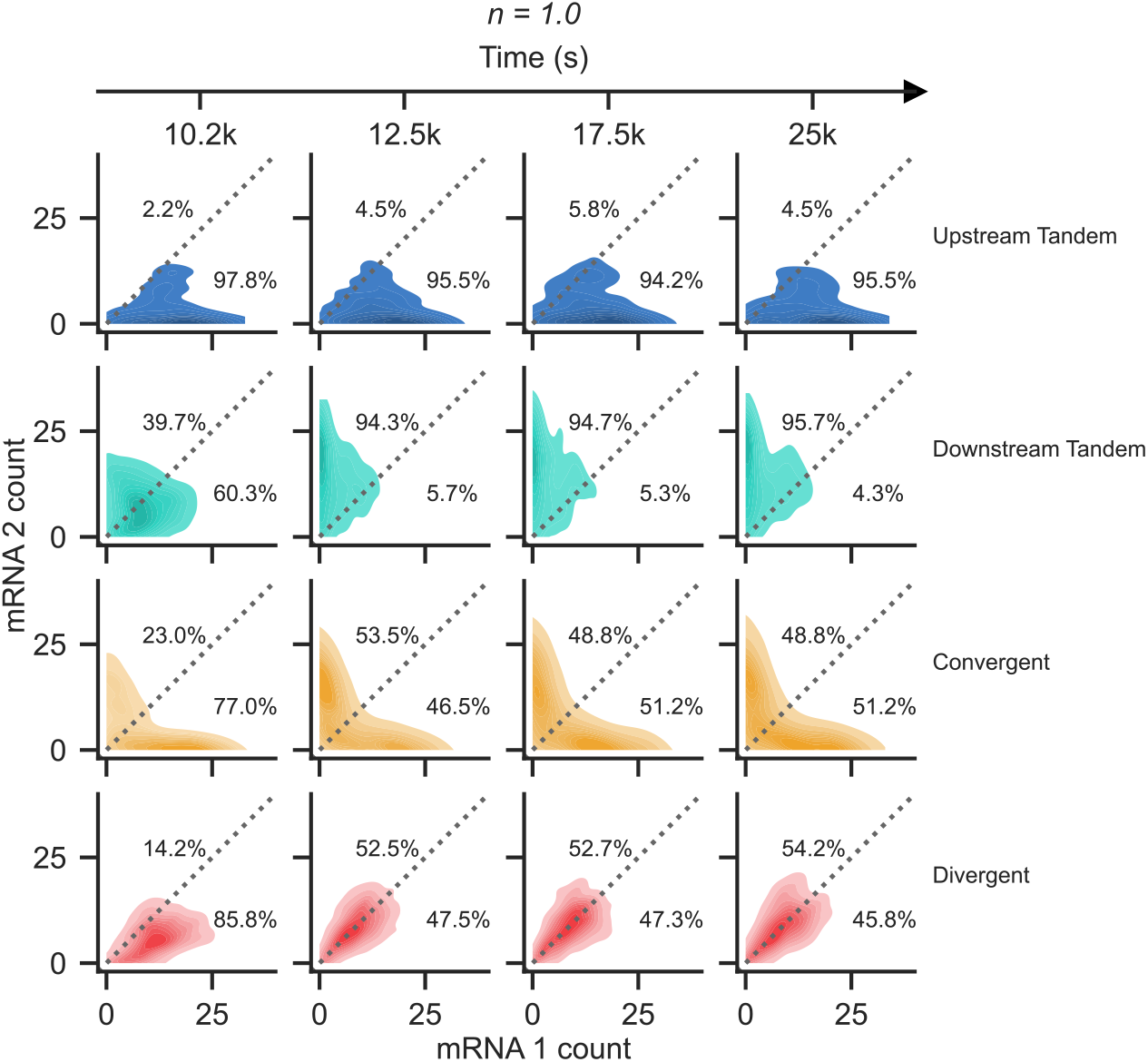
The ensemble mRNA count distributions are shown as a function of syntax at four selected time points for *n* = 1.0. Even as early as 200 seconds after induction of the second gene, we see that the tandem-downstream and divergent syntaxes are quickly approaching their equilibrium ensemble distributions.

**Figure S12:**
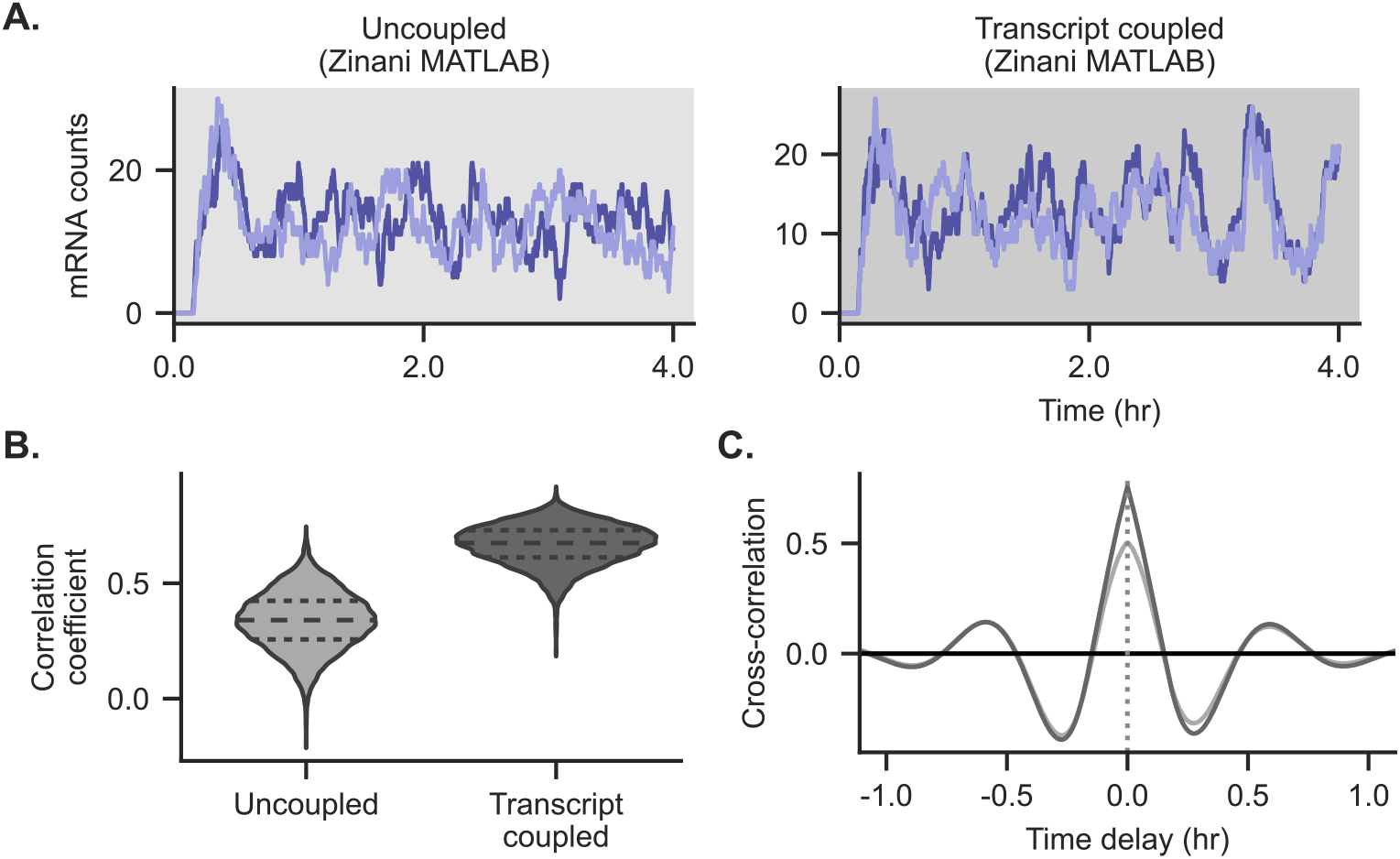
Using the authors MATLAB implementation of the model in Zinani *et al*. [47], we simulated ensembles of 10,000 runs in order to compare to our simulation work. Unlike our implementation, which does not contain extra delays not due to polymerase motion and stalling, the original model uses a fixed mRNA production delay time of 9 minutes and a fixed protein production delay time of 1 minute.a) Example runs of the original model are shown. The transcript-coupled case shows correlated but relatively aperiodic behavior.b) The ensemble correlation coefficient distributions are shown, with the transcript coupled case showing higher correlation as expected.c)The ensemble cross-correlation shows that the uncoupled and transcript coupled cases in the original model show very similar correlation structures, showing that both have similar amounts of periodicity in the original model.

**Figure S13:**
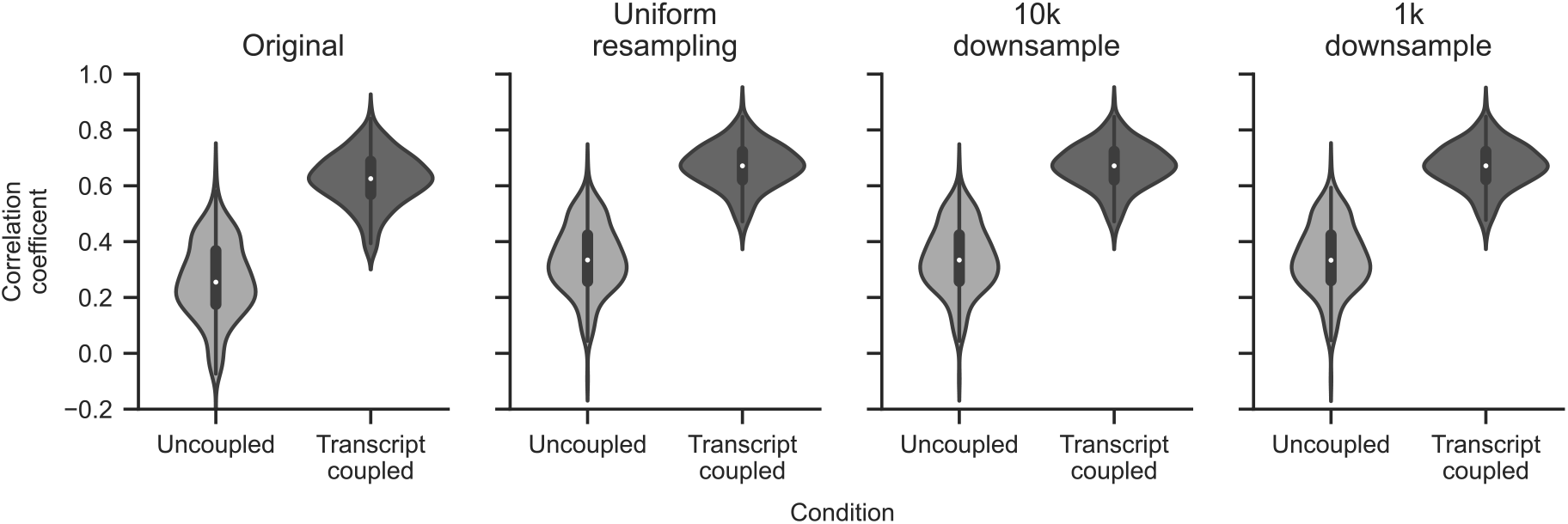
While Zinani *et al*. [47]calculate correlation coefficients using event-sampled data (far-left), we use datapoints uniformly sampled in time to calculate correlation coefficients. Using Zinani’s original MATLAB code, we computed the correlation coefficient distributions using both their original sampling method (far-left) and with the dataset resampled uniformly in time. The qualitative results between these two sampling methods remain the same, with the original sampling method slightly under-reporting *her1-her7* correlation due to its reliance on Gillespie-sampled timepoints. We further found that down-sampling the uniformly-sampled dataset did not noticeably change the correlation distributions.

**Figure S14:**
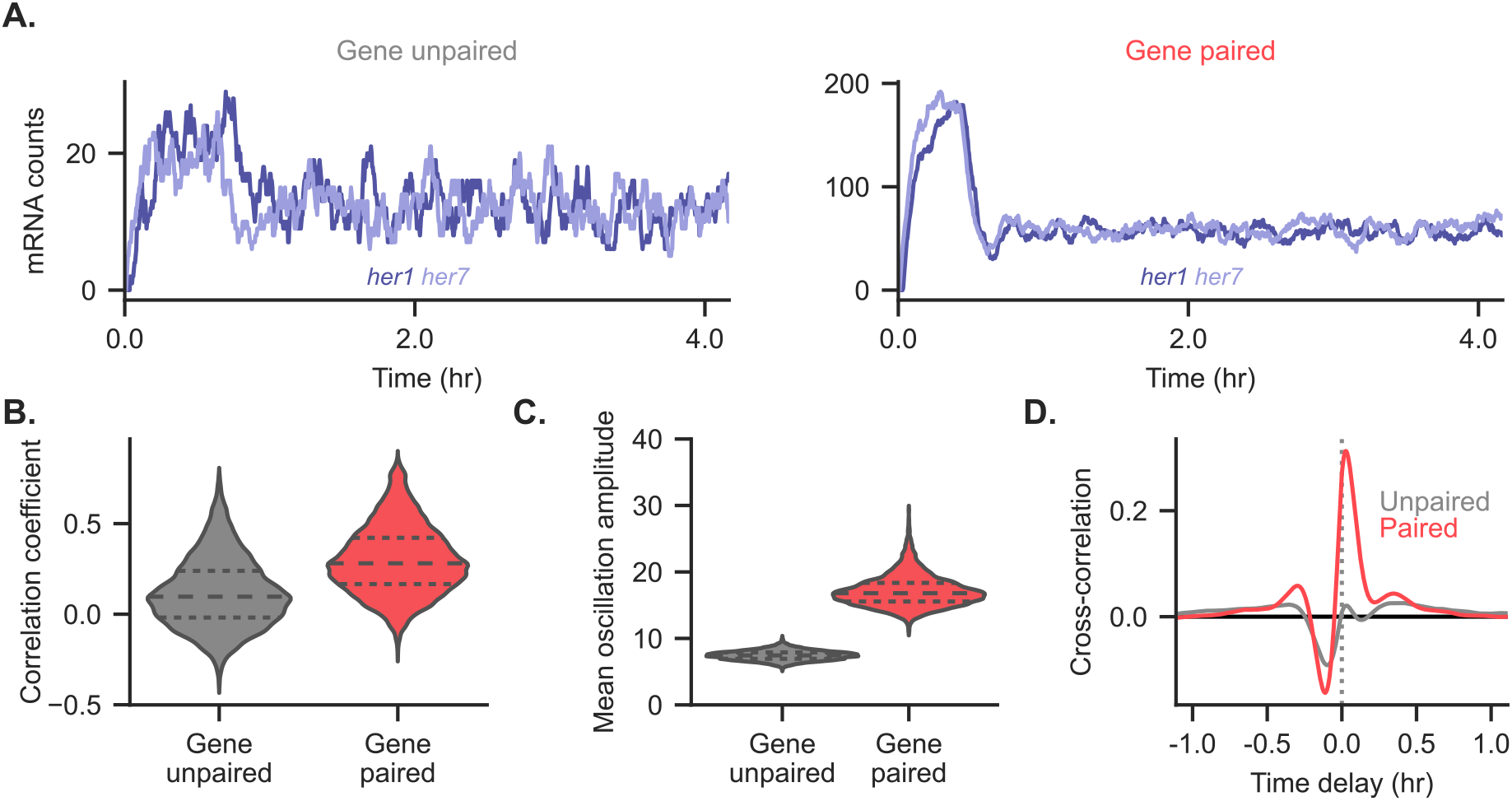
Re-simulation of the *her1-her7* system presented in fig.6 is shown for *α* = 0.02, a choice that occurs in the first regime of supercoiling-dependent initiation behavior. Strikingly, the strong correlated behavior seen in fig.6 is absent at this choice of *α*, suggesting that supercoiling-dependent initiation is an important phenomena in the synchronization of *her1* and *her7* activity. a) The number of *her1* and *her7* mRNAs are shown in the two different simulation contexts.b) In contrast to fig.6d, for *α* = 0.02, the gene paired system now shows a weaker correlated expression.c) The mean oscillation amplitude is shown for the gene unpaired and gene paired cases. The gene paired case still shows an increase in average oscillation amplitude.d) For *α* = 0.02, the biophysically-coupled case no longer shows strong periodic behavior in the cross-correlation plots, instead showing similar asymmetric cross-correlation behavior.

**Supplemental Movie 1:** An annotated example convergent simulation is shown. The top left panel shows currently loaded polymerases with growing nascent mRNAs. The bottom left panel shows the current supercoiling density plot. The right panel shows the number of mRNAs as a trace through time. The reporter gene (colored) is enabled for the length of the simulation, whereas the adjacent gene is only induced after 2.8 hours.

**Supplemental Movie 2:** An example simulation run is shown for the tandem upstream syntax. Upstream dominance, where the upstream gene has higher mean expression, is demonstrated.

**Supplemental Movie 3:** An example simulation run is shown for the tandem downstream syntax. Upstream dominance is again demonstrated.

**Supplemental Movie 4:** An example simulation run is shown for the divergent syntax. The divergent syntax demonstrates coordinated bursting.

### B Additional method details

#### B.1 Boundary conditions

Key to the our simulations is calculating the supercoiling density across the domain using eq. (1). For simulations using linear boundary conditions, we use the left and right edges as boundaries, assigning excess twist *ϕ* = 0 at both boundary locations. Then, the supercoiling density can be defined between every polymerase. The location of the boundary conditions for the simulations is described in table 2.

**Table 1:** Key dynamic constants

| Drag coefficient<br>$\eta$ (pN nm$^{\beta-1}$) | Drag exponent<br>$\beta$ | Stall torque<br>$\tau_s$ (pN nm) | Stall width<br>$\tau_w$ (pN nm) | Figures |
| --- | --- | --- | --- | --- |
| 1/20 | 1 | 12 | 3 | All except fig. S2 |
| 1/50 ↔ 1/5 | 1 | 12 | 3 | Figure S2a |
| 1/20 | 1 ↔ 3 | 12 | 3 | Figure S2b |
| 1/20 | 1 | 8 ↔ 16 | 3 | Figure S2c |
| 1/20 | 1 | 12 | 1 ↔ 5 | Figure S2d |

**Table 2:** Simulated circuit locations and distances. Locations are all given in basepairs (with 0.34 nanometers per base pair). For the given genes, the first number given in the simulated promoter location and start of the gene body, and the second number is a simulated termination site.

| Type | Boundary locations | Reporter gene | Inducible gene | Figures |
| --- | --- | --- | --- | --- |
| Upstream tandem | 0, 10000 | 3000→4000 | 6000→7000 | Figures 3b to 3e, 4 and S2 to S8 |
| Downstream tandem | 0, 10000 | 6000→7000 | 3000→4000 |  |
| Convergent | 0, 10000 | 7000→6000 | 3000→4000 |  |
| Divergent | 0, 10000 | 6000→7000 | 4000→3000 |  |
| Type | Boundary locations | Reporter gene | Inducible gene | Figures |
| Upstream tandem | 0, 8000 + $\Delta x$ | 3000→4000 | 4000 + $\Delta x$<br>→ 5000 + $\Delta x$ | Figure 3f |
| Downstream tandem | 0, 8000 + $\Delta x$ | 4000 + $\Delta x$<br>→ 5000 + $\Delta x$ | 3000→4000 | |
| Convergent | 0, 8000 + $\Delta x$ | 5000 + $\Delta x$<br>→ 4000 + $\Delta x$ | 3000→4000 | |
| Divergent | 0, 8000 + $\Delta x$ | 4000 + $\Delta x$<br>→ 5000 + $\Delta x$ | 4000→3000 | |
| Type | Boundary locations | Gene A | Gene B | Figures |
| Upstream tandem | 0, 10000 | 3500→4500 | 5500→6500 | Figures 5 and S9 to S11 |
| Downstream tandem | 0, 10000 | 5500→6500 | 3500→4500 |  |
| Convergent | 0, 10000 | 3500→4500 | 6500→5500 |  |
| Divergent | 0, 10000 | 4500→3500 | 5500→6500 |  |
| Type | Boundary locations | *her1* | *her7* | Figures |
| Gene-unpaired | 0, 10<sup>9</sup> (free ends) | 6405→6017 | (10<sup>9</sup> − 9864)<br>→ (10<sup>9</sup> − 8549) | Figures 6 and S14 |
| Gene-paired | 0, 34393 | 12422→6017 | 24529→25844 |  |

For simulations using circular boundary conditions, we must define how generated supercoiling “wraps around” the edges of the simulation. To do this, we choose an arbitrary origin, and order polymerases based on their (clockwise) position from the origin. In table 2, the first boundary location is used as this origin location and the second boundary location is the length of the circle, relative to the origin. As in the linear case, when there are zero polymerases loaded, the supercoiling density is uniformly 0. In addition, for circular boundary conditions, the supercoiling density is *also* uniformly 0; for assumed supercoiling relaxation, a single polymerase can never accumulate negative or positive supercoiling.

For two or more polymerases, we create a list of excess twists, wrapping as *ϕ_n_, ϕ*_1_,…, *ϕ_n_, ϕ*_1_. We additionally project the locations of the wrapped polymerases past the origin (e.g., the position of the wrap-around *ϕ_n_* is placed at *z*_total length_ − *z_n_*), then compute the supercoiling density as in the linear case.

#### B.2 ODE and stochastic simulation

The core ordinary differential equations were simulated using a Tsitouras’s explicit Runge-Kutte 4-5 order method [58]. Normally, one implements stochastic simulations using a time-jumping method such as Gillespie’s method. However, because the propensity of our stochastic events changes *continuously* with the continuous simulation, we need a stochastic solver that can be applied within the continuous integrator loop. Here, we used `DifferentialEquations.jl`, a performant Julia package that allows layered differential and stochastic equations. [59]

#### B.3 Computing cross-correlation

we computed the cross-correlation between the gene outputs following induction (fig. 4e), normalized by the geometric mean of the auto-correlation of these outputs. Strong negative or positive peaks evenly spaced around *τ* = 0 is a hallmark of periodic behavior, with the time offset of the peak encoding the phase offset between the signals. For two signals *f* (*t*) and *g*(*t*), the normalized cross-correlation at a certain time offset *τ* is bounded between ±1 and can be thought of as the correlation coefficient between the unshifted version of one of the signals (*f*(*t*)) and the other signal shifted in the time axis by *τ*(*g*(*t* + *τ*)). Mathematically, we use:

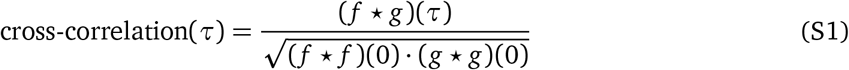

for

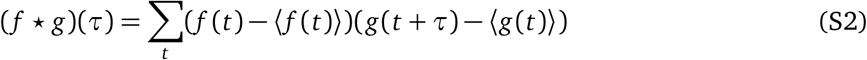

In fact, for *τ* = 0, the cross-correlation of the two signals is exactly the Pearson correlation coefficient. The shape of the cross-correlation curve can also reveal periodic and other time-dependent correlative behaviors.

#### B.4 Burst threshold choice

We calculated burst size and inter-burst time by using a burst threshold. From polymerase initiation times, we calculate the time between successive polymerase additions. Intra-addition times greater than the burst threshold form the boundaries between different expression bursts. We define the burst size to be the number of polymerases included in a burst, and the inter-burst time to be those intra-polymerase-addition times greater than the burst threshold. For the main text, we used a burst threshold of 30 seconds; specifically, this means that bursts ended if 30 seconds passed without a new polymerase being added.

In fig.S8,we reanalyze the data presented in figs. 4c and 4d for different burst thresholds. We find that using a burst threshold of twenty or sixty seconds does not significantly affect the qualitative results observed. However, using a ten second burst threshold does dramatically shift the resulting burst size and inter-burst time distributions, with the inter-burst time distribution becoming concentrated around ten seconds. This indicates that a ten-second burst time is too short and incorrectly separates bursts.

#### B.5 Choice of the half-maximum constant in the toggle-switch Hill functions

The half-max value *K*, the mRNA count at which the promoter activity is half that of *r*_0_, is chosen here to approximately match the mean steady-state expression of the steady states. The mean steady-state value is identified using simulations where only one of the toggle switch genes is enabled; this allows us to directly account for the influence of supercoiling-mediated feedback on the steady state mRNA concentration. With this choice of *K*, we ensure that the toggle switch operates in the regime of maximum sensitivity (e.g. the stable basin steady-state value is in the middle of the sigmoidal repression curve).

In fig. 5e, we tune the mRNA degradation rate, which directly impacts the mean steady-state value. If the mRNA degradation rate is doubled, we expect that the mean steady-state value should decrease to half its original value. To compare between these otherwise disparate conditions, we scaled the *K* value alongside the mRNA degradation rate, dividing by the fold increase in the mRNA degradation rate.

#### B.6 Relevant simulation parameters

### C Model design and derivation

Our model builds upon several prior papers and relevant mathematical frameworks. To ensure access and provide definitions needed for implementation of our model, we have aggregated all of these relevant equations and simulation constants together. From this foundation, we discuss how we have extended our model to examine how DNA supercoiling generates coupling between proximal genes.

#### C.1 Literature supercoiling model

Why do we care about supercoiling at all? From statistical mechanics, the thermal energy unit *k_B_T*, defined by the product of the Boltzmann constant and the temperature of the system, defines an energy accessible by fluctuations; useful chemical reactions and interactions often occur above this fluctuation energy. Converting into units relevant to supercoiling at *T* = 300 K, *k_b_T* = 4.1 pN nm, Many of the later quantities we will solve for will have magnitude around 10-100 pN nm—well above the thermal energy—-so can determining system behavior.

When we derived the supercoiling density, *σ*, in eq. (1), we defined *ϕ* to be the local excess twist. While this term is most easily identified with the local DNA excess twist, it is truly a linking number constraint that takes into account both twist, local stretching of bonds that causes more or fewer complete rotations to occur over a set distance, and writhe, where mesoscopic plectonemes store supercoiling. These two forms of supercoiling are physically interconvertable, but are not energetically identical. This energetic difference is accounted for in the underlying statistical mechanical model (fig.S1a). The statistical mechanics model depends on the following constants:

**Table 3:** Key values for the statistical mechanics model

| Variable | Value | Reference |
| --- | --- | --- |
| $k_b$, Boltzmann constant | $0.01381 \frac{\text{pN nm}}{\text{K}}$ | - |
| $\omega_0$, relaxed DNA twist frequency | $1.85 \frac{1}{\text{nm}}$ | [44], others |
| $A$, bend persistence length | 50nm | [49] |
| $C$, twist persistence length (twisted) | $95 \pm 10\text{nm}$ | [49] |
| $P$, twist persistence length (plectonemic) | $24 \pm 3\text{nm}$ | [49] |
| $f$, applied force on DNA | 1 pN | [44] |

Especially key measurable constants are the bend persistence length *A*, the twist persistence length of stretched DNA *C*, and the twist persistence length of the plectonemic state *P*. Because plectonemes convert twist into writhe, we expect *P* < *C*.

Marko defines the following rescaled versions of *P* and *C*, *p, c*:

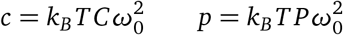

which in turn can be used to define helper constants *g, c_s_*:

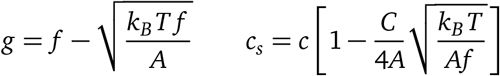

With these defined, Marko’s statistical-mechanics model calculates the torque as a function of supercoiling density (fig.S1a) [49]:

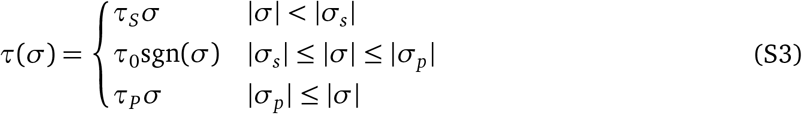

for critical supercoiling values *σ_s_, σ_p_* separating the phase regimes:

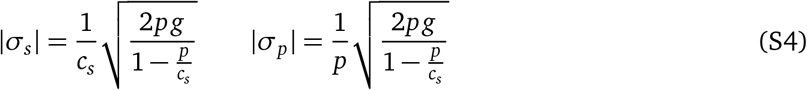

with corresponding critical torque values *τ_s_*, *τ_P_*:

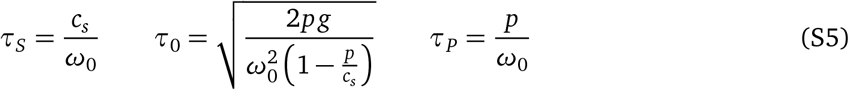

Using the torque equation eq. (S3), Marko expands the free energy of the DNA per unit length, *S*, as a power series in *σ*:

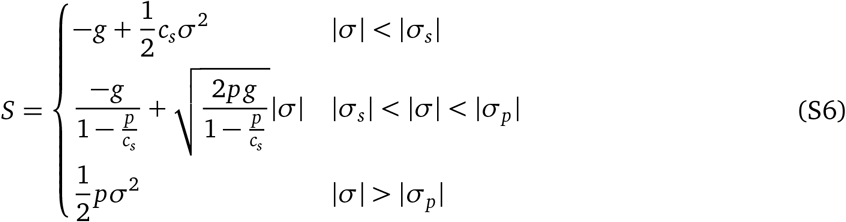

While the cubic term can also be calculated and accounts for asymmetry between under- and over-winding, Marko [49] argues that this effect is small for realistic physiological conditions.

#### C.2 Literature dynamic model

Following Sevier & Levine [44], we adopt a dynamics model for modeling the motion of polymerases over the DNA. This model depends on the following constants:

**Table 4:** Key dynamic model values

| Variable | Value | Reference |
| --- | --- | --- |
| $\chi$, DNA twist mobility | 0.01381 s pN nm | [44] |
| $\eta$, mRNA drag coefficient | 1/20 pN | [44] |
| $n$, Drag scaling exponent | 1 | [44] |
| $v_0$, Maximum polymerase velocity | $20 \frac{\text{nm}}{\text{s}}$ | [44] |
| $\tau_s$, Polymerase stall torque | 12 pN nm | [44] |
| $\Delta\tau_s$, Polymerase stall width | 3 pN nm | This work |
| $\delta$, Polymerase radius | 15 nm | [44] |

Briefly, for each polymerase, this model tracks the position *z_i_*, transcript length *x_i_*, and excess DNA twist *ϕ_i_* of each RNAP. We create a *linking number constraint* by specifying *ϕ* at any genomic location, which means that each RNAP sets an linking number constraint. We use dynamic equations to model the motion of RNAP against either fixed or free boundary conditions.

As mentioned in the main text, the dynamic equations are:

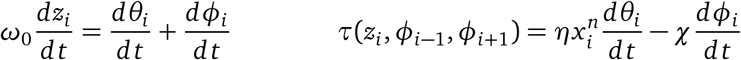

By solving the left equation for 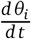, the polymerase angular velocity, and substituting it into the right equation, we get:

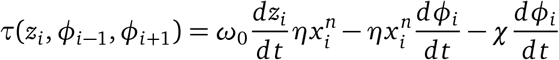

Solving for 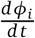, we get a dynamic equation for the differential linking number constraint:

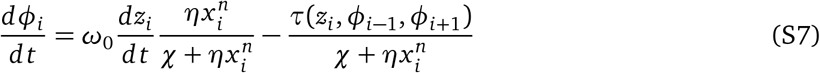

To model polymerase stalling, we use a parameterized function discussed in eq. (4) that explicitly depends on a stall torque and a torque width *τ_w_* over which stalling begins to occur.

Equations C.2 and 4 along with the definition of linear velocity 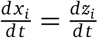 together represent a complete coupled set of ODEs that can be simulated, as long as the following constants are defined:

#### C.3 Model extension: supercoiling-dependent initiation

Reflecting on the desired properties of a model based on literature observations, we extended the models from Sevier and Marko to account for supercoiling-dependent initiation. The rest of this appendix details this process, in addition to the initial model formulation and useful simplifications.

##### C.3.1 Literature observations and modeling goals

A naked DNA bead experiment (Revyakin *et al*. [60]) found that positive and negative supercoiling affected RNAP polymerase binding rates, with negative supercoiling encouraging binding and positive supercoiling discouraging binding. When a polymerase bound, approximately 1.2 turns of DNA unwound (13 base pairs, or 4.42nm). For a typical bacterial promoter, RNAP binding occurred reversibly. The mean time between RNAP binding events was measured and related to a kinetic model. The mean time between binding events defines a promoter-on rate. Once the DNA torque reaches the constant-torque regime, the promoter-on rate becomes constant.

Ideally, our model would replicate these results, with negative supercoiling promoting RNAP binding and negative supercoiling inhibiting RNAP binding, but would only depend on the local supercoiling density. Successful phenomenological models such as El Houdaigui *et al*. [30] often use sigmodial curves that are most sensitive in the range from *σ* = −0.1 to *σ* = 0.1; ideally, our model would be sensitive in a similar regime.

##### C.3.2 Dual linking number constraint model

To model the unwinding induced by RNAP binding, we add two new linking number constraints, spaced 13 base pairs apart, and analyze the energy required to perform this unwinding. To start, all of these have specific linear locations *z* and rotations *ϕ*; we will relax this restriction later.

**Figure S15:**
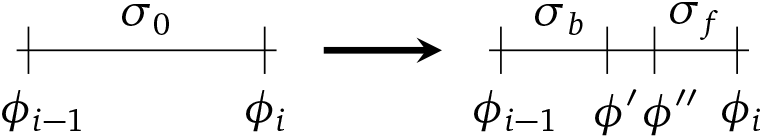
Diagram of the addition of two linking number constraints

We then put an angular restriction on the two new linking number constraints. By subtracting off the excess angle *ϕ* that the undisturbed DNA has prior to RNAP binding we can define:

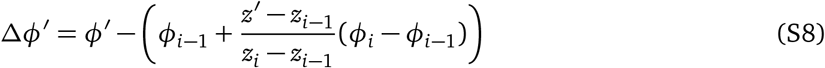

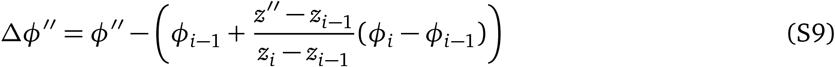

For unwinding of the region bound to the RNAP to occur, we impose two constraints that ensure that the 13 base pairs that interact with the RNAP are fully unwound:

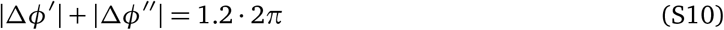

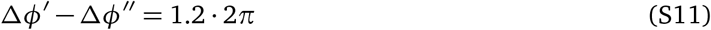

For the intermediate region, this implies that there are two different ways to unwind the DNA; either to rotate the leading linking number constraint backwards or rotate the trailing linking number constraint forwards. There are also many intermediate solutions where *both* of the bounding linking number constraints move. The first restriction is there to ensure that we use the solution that minimizes total amount of rotation needed.

Given a complete energy model that related *τ* over the complete range of unwinding Δ*ϕ′*, Δ*ϕ″* values, we could directly write the binding energy as

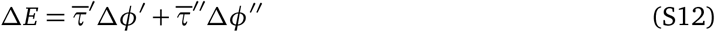

and minimize the energy cost with respect to Δ*ϕ′* or Δ*ϕ″*, giving a unique solution for any specific geometry. As Marko’s statistical mechanical model is likely not valid in the limit of complete unwinding, we instead move forward with this model by estimating the energy cost in both the “exterior” region and within the “interior”, 13-bp region.

##### C.3.3 Energy estimation in the exterior region

If we assume that the energy surface is locally linear (e.g. in the exterior region, there is a small change in supercoiling density), then we can relate the instantaneous torque to the change in energy:

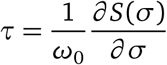

for *S*, the energy per unit length. This means that:

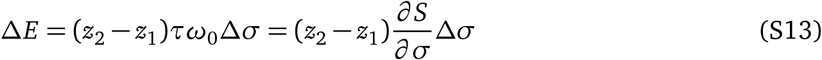

By expanding the definition of Δ*σ* for an arbitrary region bounded by two linking number constraints, we can actually recover the normal energy-torque definition:

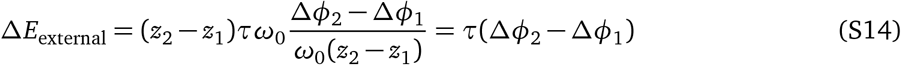

Note that this implies that the energy cost of linking number constraint rotation is independent of the region width; under a locally linear assumption, the energy cost depends only on the local instantaneous torque and the rotation angle. When is this valid? The change in supercoiling density in the exterior region is relatively low, *O*(0.05) as long as the region of interest is at least 250*bp* away from the nearest linking number constraint on one of its sides. For promoter regions analyzed here, space between gene bodies ensures that this assumption holds.

##### C.3.4 Energy estimation in the interior region (local melting)

We can generally write the energy it takes to unwind the DNA in the interior region as an integral over the linear free energy density, *S*:

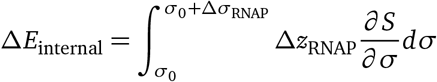

Given the well-defined structure of the RNAP-DNA complex and the fact that under all realistic physiological conditions Δ*σ*_RNAP_ ≫ *σ*_0_, we assume that the end-state energy is a constant, but unknown *S*_unwound_:

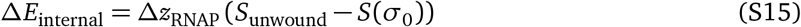

While *S*_unwound_ is unknown, it is a constant with respect to all promoters, so this energetic term is implicitly already included in any promoter base rate. This means that the external energy depends only on the free energy density, evaluated at the local, initial supercoiling density *σ*_0_:

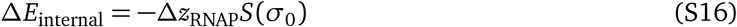

##### C.3.5 Order of magnitude energy analysis

Combining the results of eqs. (S14) and (S16) and plugging in eq. (S11), we have an estimated binding energy:

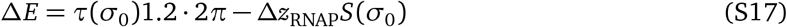

When we plot this energetic term in fig.S1b, we see that the internal energetic term is minimal when compared to the external energetic term (local melting) at the supercoiling densities considered here, so we use the simplified form

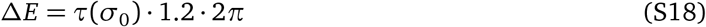

as presented in eq. (5).

## Notes

https://github.com/GallowayLabMIT/tangles_model

### Competing Interest Statement

The authors have declared no competing interest.

### Summary of Updates

Fixed citation format in references (removed URL download dates) and added supplemental movie captions.

